# Generation of Valvular Interstitial Cells from Human Pluripotent Stem Cells

**DOI:** 10.1101/2025.09.17.676963

**Authors:** Amine Mazine, Alexander A. Mikryukov, Ian Fernandes, Clifford Z. Liu, Soheil Jahangiri, Marcy Martin, Eric K. N. Gähwiler, Melanie Generali, Juliana Gomez, Neda Latifi, Yifei Miao, Yu Liu, Michael A Laflamme, Craig A Simmons, Simon P. Hoerstrup, Maximilian Y. Emmert, Bruce D. Gelb, Mingxia Gu, Gordon M. Keller

**Affiliations:** McEwen Stem Cell Institute, University Health Network, Toronto, ON, Canada; Institute of Biomedical Engineering, University of Toronto, Toronto, ON, Canada; Division of Cardiac Surgery, Department of Surgery, University of Toronto, Toronto, ON, Canada; Mindich Child Health and Development Institute, Icahn School of Medicine at Mount Sinai, New York, NY, USA; Institute for Regenerative Medicine, University of Zurich, Zurich, Switzerland; Department of Medical Engineering, University of South Florida, Tampa, FL, USA; Division of Pulmonary Biology, Perinatal Institute, Cincinnati Children’s Hospital Medical Center, Department of Pediatrics, University of Cincinnati College of Medicine, Cincinnati, OH, USA; Center for Stem Cell and Organoid Medicine (CuSTOM), Cincinnati Children’s Hospital Medical Center, Department of Pediatrics, University of Cincinnati College of Medicine, Cincinnati, OH, USA; Division of Developmental Biology, Cincinnati Children’s Hospital Medical Center, Department of Pediatrics, University of Cincinnati College of Medicine, Cincinnati, OH, USA; Cardiovascular Institute, Stanford School of Medicine, Stanford, CA, USA; Department of Medicine, Division of Cardiovascular Medicine, Stanford School of Medicine, Stanford, CA, USA; Department of Cardiothoracic and Vascular Surgery, Deutsches Herzzentrum der Charite (DHZC), Berlin, Germany; Departments of Pediatrics and Genetics and Genomic Sciences, Icahn School of Medicine at Mount Sinai, New York, NY, USA; Princess Margaret Cancer Centre, University Health Network, Toronto, ON, Canada; Department of Medical Biophysics, University of Toronto, Toronto, ON M5G1L7, Canada

**Author notes:** Correspondence (A.M.), (G.M.K.). contributed equally (Amine Mazine, Alexander A. Mikryukov).

## Abstract

Heart valves are living structures whose sophisticated functions are mediated by a specialized population of mesenchymal cells known as valvular interstitial cells (VICs). Given their central role in valve homeostasis, VICs represent a promising cell population for studying heart valve diseases and developing novel therapies to treat them. Here, we describe a strategy for generating VICs from human pluripotent stem cells (hPSCs) by stage-specific manipulation of developmental signalling pathways. Our results demonstrate that hPSC-derived VICs show a high transcriptional similarity to primary human fetal VICs and can secrete key proteins of the valve extracellular matrix. We further investigate the heterogeneity of hPSC-derived VICs and identify two major subpopulations with distinct molecular and functional properties, mirroring the cellular diversity observed *in vivo*. Finally, we utilize an *in vitro* model of Noonan syndrome to demonstrate that hPSC-derived VICs can accurately recapitulate key aspects of valve disease. Collectively, these findings provide a reproducible method for the scaled generation of *bona fide* hPSC-derived VICs and establish their utility in disease modelling and tissue engineering applications.

**Clinical Perspective:** *What is new?:*

- We established a robust platform to generate *bona fide* valvular interstitial cells (VICs) from human pluripotent stem cells (hPSCs), recapitulating native VIC identity and function.
- We delineated signaling pathways that promote the development of two distinct VIC subsets and identified a surface marker to distinguish between them.

*What are the clinical implications?:*

- A renewable, human-specific source of VICs enables precise mechanistic studies of valve development and disease that are not possible with limited surgical specimens or animal models.
- This platform creates opportunities for therapeutic applications, including drug discovery and tissue engineering approaches for valve repair.

---

Heart valves are anatomically specialized living structures that allow unidirectional blood flow through the heart chambers and into the great vessels. They are composed of thin (∼500 μm) pliable leaflets lined with a single layer of valvular endothelial cells (VECs) and populated with unique mesenchymal cells known as valvular interstitial cells (VICs), which can be distinguished from other cardiac mesenchymal cells by their elevated expression of a select set of genes that include *PRRX2*, *TIMP3*, and *ITGA2* ^1^. VICs play a crucial role in ensuring the proper functioning and longevity of heart valves by secreting and maintaining the highly compartmentalized extracellular matrix (ECM) of the valve leaflet ^2^. This ECM comprises multiple layers, including a central *spongiosa* layer, rich in proteoglycans and glycosaminoglycans, which imparts compressibility and flexibility to the leaflets, and a dense collagen-rich *fibrosa* layer, which provides stiffness and tensile strength ^3^. Single-cell RNA sequencing (scRNA-seq) analyses of neonatal murine heart valves have identified two distinct subpopulations of VICs, one that expresses genes involved in collagen fibril organization and ECM maturation (predominantly located in the *fibrosa* layer) and a second that expresses glycosaminoglycan-related genes (predominantly in the *spongiosa* layer) ^4^. Given their central role in valve homeostasis, VICs represent a promising cell population for studying heart valve diseases and developing novel therapies to treat them.

Valvular heart disease can affect patients of all ages and is an important and growing cause of global cardiovascular morbidity and mortality ^5^. Irrespective of etiology, advanced valve disease is characterized by structural damage that impairs proper valve function. As only a small fraction of native valves with structural pathology can be effectively repaired, the mainstay of treatment remains prosthetic valve replacement. However, prosthetic valves are acellular non-living substitutes; they cannot recapitulate the sophisticated functions of a normal heart valve and are prone to degeneration, thrombosis, and infection. As a result, children and non-elderly adults who undergo prosthetic valve replacement have a curtailed life expectancy compared with the age- and sex-matched general population ^6^.

A tissue-engineered living heart valve substitute, populated by appropriate cell types that faithfully replicate the native heart valve’s functions, represents a promising alternative to currently available prostheses, particularly in young patients. To achieve this, the tissue-engineered valve should possess the ability to grow alongside the patient, maintain its structural integrity and functionality throughout the patient’s lifetime, and minimize inflammation and adverse remodelling. However, developing such a living substitute is contingent on our ability to access VICs in sufficient quantities. While numerous approaches involving various biomaterials, maturation techniques, and cell sources have been explored ^7^, none has utilized *bona fide* human VICs, as access to these cells is limited by the scarcity of organ donors and the lack of appropriate conditions to expand them *in vitro*. Human pluripotent stem cells (hPSCs) represent an alternative and potentially inexhaustible source of VICs. As with other cell types ^8–11^, the successful derivation of functional VICs from hPSCs will depend on our ability to translate critical aspects of their embryonic development to the differentiation cultures. Lineage tracing studies in model organisms have demonstrated that most VICs arise from a subset of endocardial cells that line the endocardial cushions ^12, 13^. These cushion endocardial cells exhibit molecular and functional properties that are distinct from those of chamber endocardium ^14–16^. Through the coordinated interaction of different signalling pathways, including BMP, TGFβ, and Wnt/β-catenin, these cells undergo an endothelial-to-mesenchymal transition (EndoMT) to form cushion mesenchymal cells that eventually give rise to VICs ^17^.

Our group has previously developed a protocol to generate endocardial cells from hPSCs ^9^. In the present study, we identify a subpopulation of these hPSC-derived endocardial cells, characterized by the co-expression of CD31, NKX2-5, and PDGFRβ, that can undergo EndoMT and give rise to cells that display properties of VICs. We demonstrate that these VIC-like cells show a high level of transcriptional similarity to primary human fetal VICs and can secrete key proteins of the valve ECM. We further investigate the heterogeneity of hPSC-derived VICs and identify two major subpopulations with distinct molecular and functional properties, mirroring the cellular diversity seen *in vivo*. Finally, we utilize an *in vitro* Noonan syndrome iPSC model to demonstrate that hPSC-derived VICs can accurately recapitulate key aspects of valve disease.

## Methods

Detailed descriptions of cell culture, differentiation protocols, flow cytometry, immunohistochemistry, tissue engineering, and sequencing methods are available in the Supplemental Methods.

### Institutional approval

Studies involving human fetal tissue were conducted in collaboration with the University of Washington Birth Defects Research Lab (BDRL, IRB STUDY00000380), University of Southern California and Children’s Hospital Los Angeles, and Cincinnati Children’s Research Foundation (CCRF, IRB 2011-2856). All samples were de-identified and collected with the signed informed consent of parents in accordance with institutional, state, and federal regulations. The only clinical information collected was gestational age and the presence of any maternal or fetal diagnoses. The work presented herein was classified as exempt from further IRB review because of the de-identified nature of the valve samples.

### Data availability

Single-cell RNA sequencing data generated in this study have been deposited in the Gene Expression Omnibus (GEO) under accession numbers **GSE228638**, **GSE151186**, and **GSE234382**. Additional data supporting the findings of this study are available from the corresponding author upon reasonable request.

### Statistical analysis

All data are represented as mean ± standard error of mean (SEM). Sample sizes (n) represent biological replicates of differentiation experiments. No statistical method was used to predetermine the samples size. Due to the nature of the experiments, randomization was not performed and the investigators were not blinded. Statistical significance was determined by using Student’s t test (paired, two-tailed) and one-way or two-way analysis of variance (ANOVA) with post-hoc testing using Tukey correction for multiple comparisons in GraphPad Prism 7 software. Results were considered to be significant at p < 0.05 (*), p < 0.005 (**), p < 0.0005 (***). All statistical parameters are reported in the respective figures and figure legends.

## Results

### PDGFRβ expression defines subpopulations of endocardial cells

For these studies, hPSCs were differentiated into endocardial-like cells using our previously published protocol^9^ that includes the induction of cardiogenic mesoderm with specific concentrations of activin A and BMP4 (3 ng/mL activin A, 5 ng/mL BMP4), and the specification of this mesoderm to an endocardial fate by the addition of bFGF and BMP10 (**Fig 1a**). To further characterize the heterogeneity of the hPSC-derived endocardial NKX2-5^+^ population, we re-analyzed our previous scRNA-seq data^9^, focusing on genes that encode surface markers. From this analysis, we identified a subset of cells that express *PDGFRβ*, the gene that encodes the PDGFRβ receptor found on mesenchymal cells^18^. Of the 1,063 NKX2-5^+^ cells profiled, 265 (24.9%) were categorized as *CD31^+^PDGFRβ^+^*, while 498 (46.8%) were identified as *CD31^+^PDGFRβ^-^* (**Fig 1b–d**). Gene set enrichment analysis (GSEA) showed enrichment of genes associated with epithelial-to-mesenchymal transition, mesenchyme development and endocardial cushion development in the *CD31^+^PDGFRβ**^+^*** cells compared to the *CD31^+^PDGFRβ**^-^***cells (**Fig 1d**).

**Figure 1.**
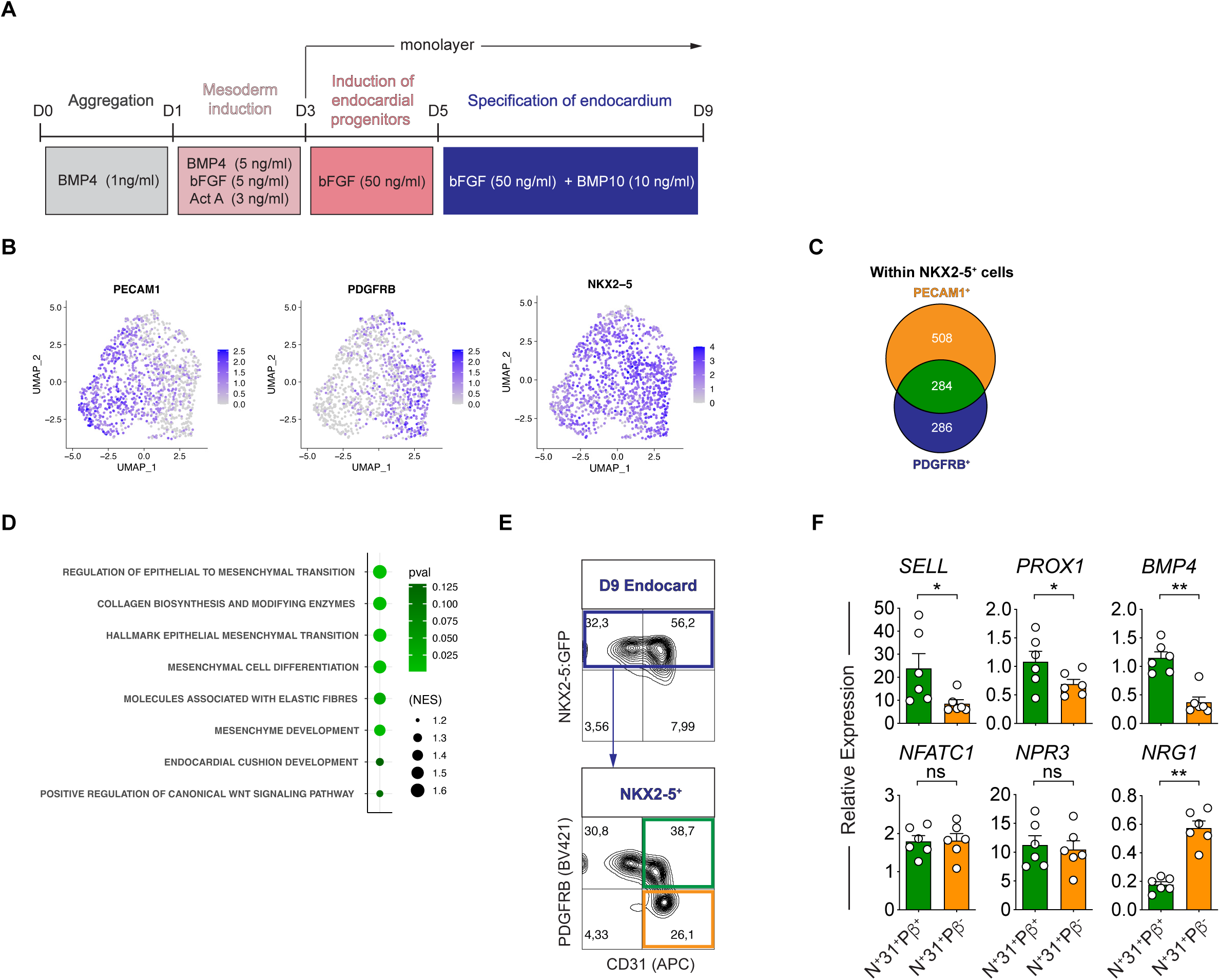
hPSC-derived PDGFRβ^+^ endocardial cells express VEC markers. **(A)** Schematic of the protocol used for the generation and isolation of hESC-derived NKX2-5^+^CD31^+^ endocardial-like cells. **(B)** UMAP plots of Day 9 hPSC-derived bulk endocardial populations generated with the protocol in (A) showing expression patterns of pan-endothelial (*PECAM1*), cardiac progenitor (*NKX2-5*) and mesenchymal (*PDGFRβ*) genes. **(C)** Venn diagram of number of *NKX2-5*^+^ cells from (B) expressing *PECAM1* (*CD31*) and (*PDGFRβ*). **(D)** GSEA showing pathways that are enriched in *PDGFRβ*^+^ compared to *PDGFRβ^-^ NKX2-5^+^CD31^+^* endocardial subpopulations from (C). (**E)** Representative flow cytometric analyses of NKX2-5/CD31 expression in endocardial cells generated with the protocol in (A) and expression of PDGFRβ on a subpopulation of NKX2-5^+^CD31^+^ endocardial-like cells. **(F)** RT-qPCR analysis of the expression levels of valvular endothelial cell-specific genes (*SELL1*, *NFATC1*, *PROX1*, *BMP4*) and chamber-specific endocardial genes (*NRG1*, *NPR3*) in PDGFRβ positive and negative NKX2-5^+^CD31^+^ endocardial subpopulations (n = 6, paired t-test *p < 0.05, ** p < 0.01, *** p < 0.005; error bars ± SEM). For all RT-qPCR analyses, expression values were normalized to the housekeeping gene *TBP*. Error bars represent SEM. *D*, day.

Flow cytometric analyses confirmed this heterogeneity and identified distinct CD31^+^PDGFRβ^+^ and CD31^+^PDGFRβ^-^ fractions within the NKX2-5^+^ endocardial population (**Fig 1e**). qRT-PCR analyses of FACS-isolated fractions showed higher levels of expression of genes associated with VIC progenitors including *SELL*, *PROX1* and *BMP4* ^1, 19^ in the CD31^+^PDGFRβ^+^ cells than in the CD31^+^PDGFRβ^-^ cells (**Fig 1f**). In contrast, the PDGFRβ^-^ endocardial cells expressed higher levels of *NRG1*, a marker of chamber endocardium known to regulate the process of ventricular trabeculation^20^. Together, these findings clearly demonstrate heterogeneity within the NKX2-5^+^CD31^+^endocardial population and raise the interesting possibility that the NKX2-5^+^CD31^+^PDGFRβ**^+^**cells represent VIC-like progenitors, poised to undergo a mesenchymal transition.

### hPSC-derived PDGFRβ^+^ endocardial cells display responsiveness to ATP

Endocardial cells in different regions of the heart are exposed to different biomechanical forces. Fukui *et al*. recently identified an extracellular ATP-dependent P2 purinergic receptor pathway that triggers calcium oscillations in response to mechanical forces ^14^. These calcium oscillations result in the nuclear translocation and activation of NFATc1, a calcium-sensitive transcription factor known to modulate EndoMT and subsequent heart valve morphogenesis (**Fig 2a**). Importantly, this group demonstrated that this signalling pathway was only active in the endocardial cells lining the valve-forming region, which contains the VIC progenitors. To determine if the NKX2-5^+^CD31^+^PDGFRβ**^+^**subpopulation of endocardial cells exhibits this characteristic response, isolated cells were incubated with the fluorescent calcium indicator Fluo-4 and subsequently stimulated with ATP (**Fig 2b**). Optical mapping of intracellular calcium transients was used to measure the cells’ response to ATP. The response of the NKX2-5^+^CD31^+^PDGFRβ**^+^**endocardial cells was compared to that of hPSC-derived CD31^+^PDGFRβ^-^ endocardial cells, hPSC-derived vascular endothelial cells, and primary human umbilical vein endothelial cells (HUVECs). Following the addition of ATP, the NKX2-5^+^CD31^+^PDGFRβ**^+^** endocardial cells displayed calcium transients with a higher amplitude and a larger area under the curve than observed in the CD31^+^PDGFRβ^-^ endocardial cells or in the other endothelial populations (**Fig 2c–e**). This response was abrogated when the cells were treated with the P2 purinergic antagonist PPADS (**Fig 2f–h**), demonstrating that the calcium transients observed in response to changes in extracellular ATP levels are modulated, at least in part, by P2 purinergic channels. These findings further support the interpretation that the CD31^+^PDGFRβ**^+^**endocardial cells are distinct from the CD31^+^PDGFRβ^-^ cells and display properties of the valve-forming region of the endocardium.

**Figure 2.**
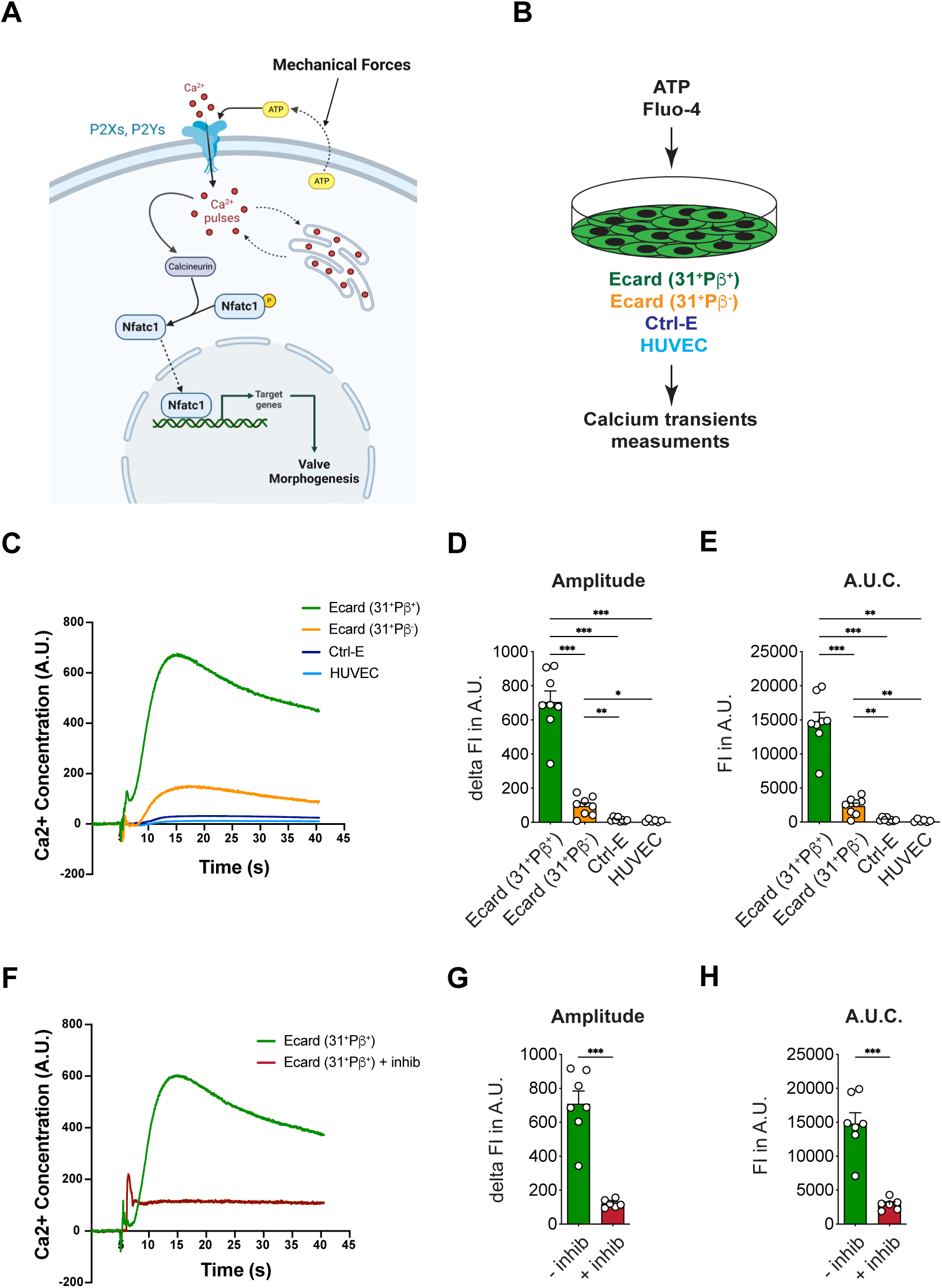
hPSC-derived PDGFRβ^+^ endocardial cells display VEC-like responsiveness to ATP. **(A)** Schematic of ATP dependent activation of NFATC signaling by Ca^2+^ influx in valve endocardial cells. **(B)** Schematic of experimental design used for optical mapping of intracellular calcium transients in various endocardial and endothelial populations. **(C)** Time course of calcium transient concentration in various endocardial and endothelial populations upon stimulation with ATP. **(D, E)** Quantification of the amplitude (D) and area under curve (E) of calcium transients measurements in (C). (n = 8, one-way ANOVA with Tukey’s multiple comparisons test, *p < 0.05, **p < 0.01,***p < 0.001; error bars ± SEM). **(F)** Time course of calcium transient concentration in NKX2-5^+^CD31^+^PDGFRβ^+^ endocardial cells upon stimulation with ATP in the presence or absence of the P2 purinergic antagonist PPADS. **(G, H)** Quantification of the amplitude (G) and area under curve (H) of calcium transient measurements in (F). (n = 7, Student’s t-test *p < 0.05, ** p < 0.01, *** p < 0.001; error bars ± SEM). Error bars represent SEM. *A.U.C.*, area under curve; *Ctrl-E*, hPSC-derived vascular endothelial cells; *ns*, non-significant.

### hPSC-derived PDGFRβ^+^ endocardial cells can undergo EndoMT and generate mesenchymal cells

The ability to undergo EndoMT and generate VICs is a defining feature of the subpopulation of endocardial cells that line the endocardial cushions ^21^. We have previously shown that the hPSC-derived endocardial population can undergo EndoMT and generate mesenchymal progeny that expresses genes associated with human fetal VICs ^9^. To determine which CD31^+^ subpopulation displays this potential, we isolated both the CD31^+^PDGFRβ**^+^** and CD31^+^PDGFRβ**^-^**cells by FACS and cultured them in the presence of BMP4 and TGFβ2, two key inducers of valve EndoMT *in vivo* (**Fig 3a**). Following eight days of culture, the CD31^+^PDGFRβ**^+^,** but not the CD31^+^PDGFRβ**^-^**, endocardial cells generated a distinct CD31^-^PDGFRβ^+^ mesenchymal population (**Fig 3b–e**). These findings show that the capacity to undergo EndoMT tracks with the CD31^+^PDGFRβ**^+^** endocardial subpopulation, consistent with the interpretation that these cells represent the *in vitro* equivalent of the endocardium that lines the endocardial cushions and gives rise to VICs in the fetal heart.

**Figure 3.**
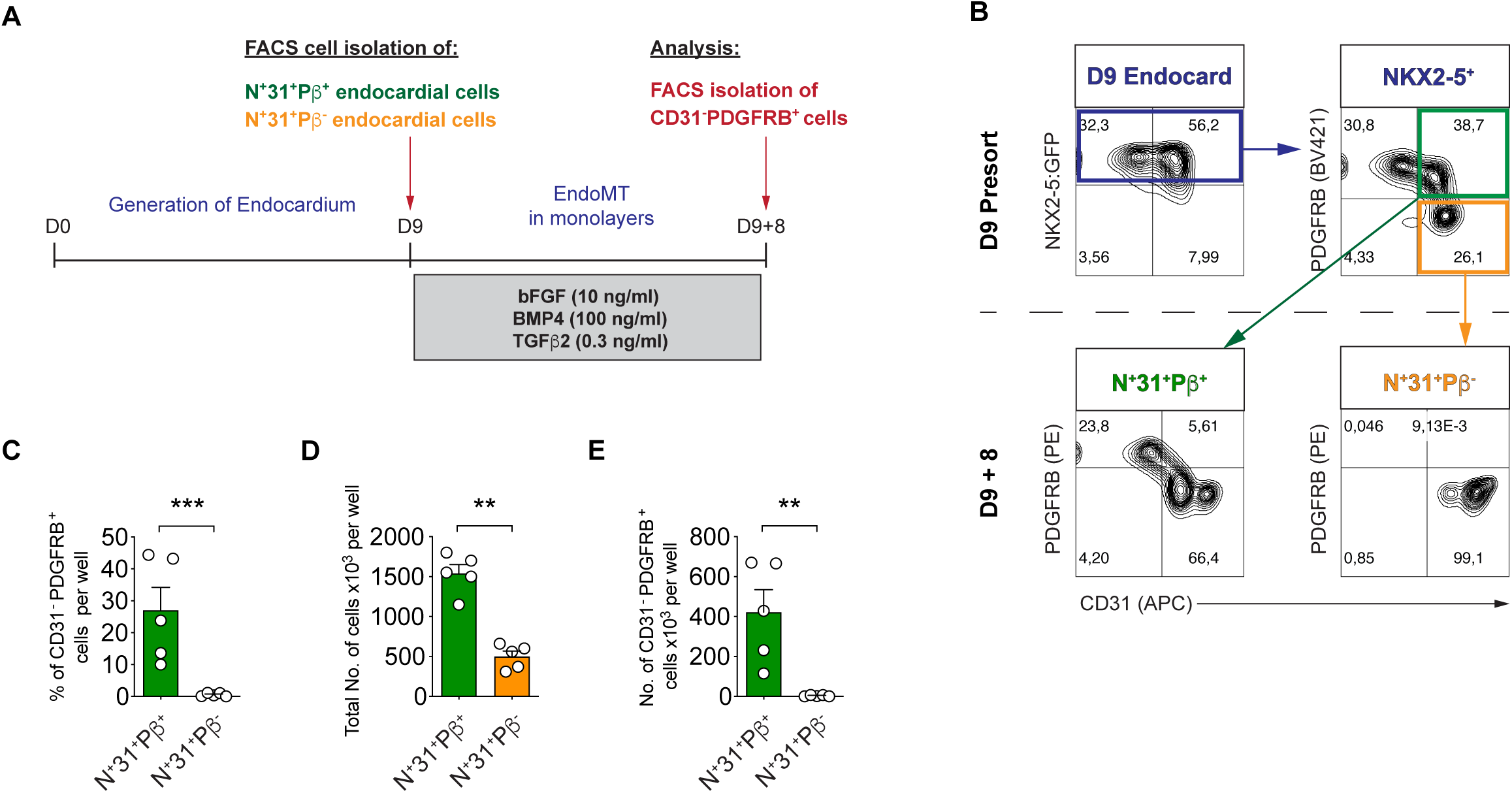
hPSC-derived PDGFRβ+ endocardial cells can undergo EndoMT. **(A)** Schematic of the protocol used to generate and isolate hPSC-derived CD31^-^PDGFRβ^+^ mesenchymal cells from PDGFRβ positive and negative NKX2-5^+^CD31^+^ endocardial populations. **(B)** Representative flow cytometric analyses of NKX2-5, CD31, and PDGFRβ expression on the day 9 endocardial population and of PDGFRβ and CD31 expression on the mesenchymal cells generated from the isolated PDGFRβ positive and negative NKX2-5^+^CD31^+^ endocardial cells. **(C-E)** Quantification of the frequency of CD31^-^PDGFRβ^+^ cells (C), of the total number of cells (D), and the total number of CD31^-^PDGFRβ^+^ cells (E) derived from PDGFRβ positive and negative NKX2-5^+^CD31^+^ endocardial cells following the protocol described in (A). (n = 5, paired t-test * p < 0.05, ** p < 0.01). Error bars represent SEM. *D*, day; *EndoMT*, endothelial-to-mesenchymal transition; *FACS*, fluorescence-activated cell sorting.

To determine whether mesenchymal cells derived from the CD31⁺PDGFRβ⁺ endocardial population exhibit mechanosensitive activation similar to primary VICs, we cultured them on substrates of differing stiffness (5 kPa versus 1.72 MPa) and assessed α-smooth muscle actin (ACTA2) expression by immunocytochemistry. Cells cultured on the stiff substrate exhibited a pronounced increase in ACTA2 expression and well-defined stress fibers, consistent with myofibroblastic activation (**Supplementary Fig. 1**). In contrast, cells on the soft substrate displayed minimal ACTA2 expression. These results demonstrate that mesenchymal cells derived from the CD31⁺PDGFRβ⁺ endocardial population are capable of stiffness-induced activation, recapitulating a key feature of native VICs.

### Wnt signaling enhances the generation of VIC-like cells from hPSC-derived endocardial cells

While the above studies demonstrated that the hPSC-derived CD31^+^PDGFRβ^+^ endocardial cells can undergo EndoMT, the process is not efficient as the resulting population is mixed and consists of only 25-30% CD31^-^PDGFRβ^+^ mesenchymal cells. To improve the efficiency of EndoMT, we next investigated the effects of Wnt signalling on this process, as previous studies have shown that this pathway plays a pivotal role in the induction of valve EndoMT in model organisms *in vivo* ^22^. For these studies, we evaluated the effect of Wnt activation on D9 endocardial cells by the addition of the GSK-3 inhibitor CHIR99021 (CHIR) in the presence or absence of BMP4 and bFGF for four days prior to the addition of TGFβ2 for an additional eight days (VIC specification phase) (**Fig 4a**). The rationale was to expand the VIC progenitor population before the induction of EndoMT. CHIR was paired with the TGFβ inhibitor SB-431542 (SB) to eliminate potential endogenous TGFβ signalling during the expansion phase. Treatment with CHIR and SB for this four-day period did not significantly impact the ratio of CD31^+^PDGFRβ^-^ : CD31^-^PDGFRβ^+^ cells (**Fig 4b**). This ratio was primarily driven by the presence of BMP4 and bFGF signalling. Eight days following the addition of TGFβ2, the populations generated under the four different conditions all contained greater than 90% CD31^-^ PDGFRβ^+^ cells. However, the number of these cells differed significantly. The D4 population expanded in the presence of CHIR, SB, BMP4, and bFGF generated the highest number of CD31^-^PDGFRβ^+^ cells, significantly more than the populations treated with either CHIR/SB or BMP4/bFGF alone (**Fig 4c**). Compared with our original EndoMT protocol (**Fig 3a)**, this modified protocol generates 4.5-fold higher frequency and 5-fold higher numbers of CD31^-^ PDGFRβ^+^ cells from the D9 endocardial cells (**Fig 4d–g**). Importantly, qRT-PCR analyses showed that the CD31^-^PDGFRβ^+^ mesenchymal cells generated with this modified protocol expressed higher levels of the EndoMT transcription factors *SOX9* and *MSX2*, and the VIC lineage markers *NR4A2*, *PRRX2*, and *TIMP3* (**Fig 4h**) than those generated using the original protocol (**Fig 3a)**. Collectively, these findings show that activation of the Wnt signalling pathway along with TGFβ inhibition prior to treatment with TGFβ2 significantly increased the number of VIC-like cells generated from D9 endocardial cells.

**Figure 4.**
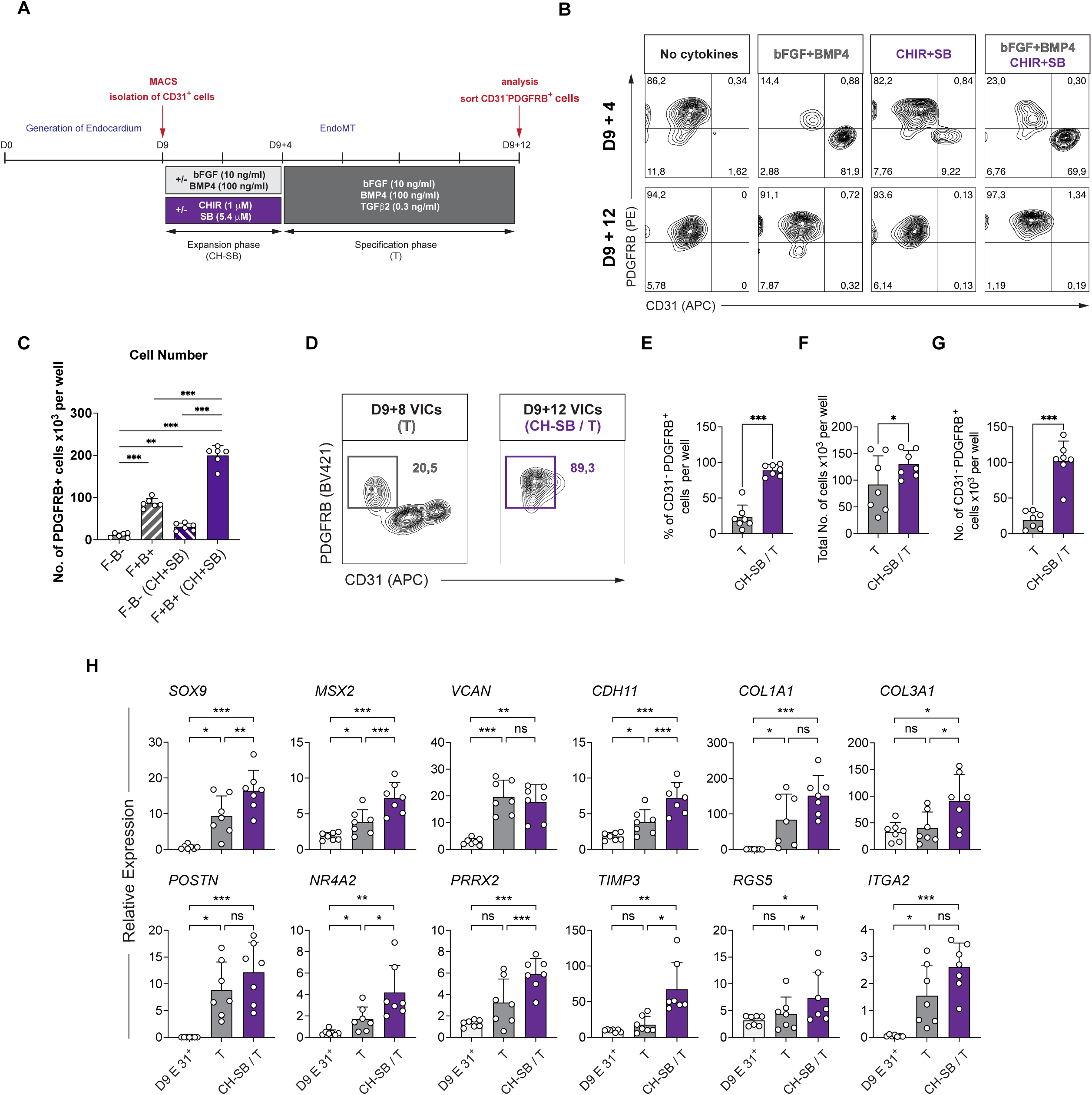
Effect of Wnt signaling on generation of VIC-like cells from hPSC-derived endocardial cells. **(A)** Schematic of the experimental strategy used to analyze the effects Wnt signaling on generation of hPSC-derived VICs. The new protocol comprises an expansion phase of 4 days followed by a specification phase of 8 days. **(B)** Representative flow cytometric analyses of CD31 and PDGFRβ expression on VICs at Day 9+4 and Day 9+12 of culture following manipulation of Wnt, BMP, FGF and TGFβ signaling during the expansion phase as indicated. **(C)** Quantification of the total number of CD31^-^PDGFRβ^+^ hPSC-derived VICs at Day 9+12 of the protocol in (A). (n = 6, one-way ANOVA with Tukey’s multiple comparisons test, *p < 0.05, **p < 0.01,***p < 0.001; error bars ± SEM). **(D)** Representative flow cytometric analyses of CD31 and PDGFRβ expression on VICs generated with the old (T) protocol (Figure 3A) and the new (CH-SB/T) protocol (Figure 4A). **(E-G)** Quantification of the frequency of CD31^-^ PDGFRβ^+^ cells (E), total cell numbers (F) and total number of CD31^-^PDGFRβ^+^ cells (G) generated with the old and new protocols in (D)(n = 7, paired t-test *p < 0.05, **p < 0.01, ***p < 0.005). **(H)** RT-qPCR analysis of the expression levels of general mesenchymal genes (*SOX9*, *MSX2*, *VCAN*, *CDH11*, *COL1A1*, *COL3A1*, *POSTN*) and VIC-specific genes (*NR4A2*, *PRRX2*, *TIMP3*, *RGS5*, *ITGA2*) in CD31^-^PDGFRβ^+^ cells generated with the old and new VIC protocols in (D). (n = 7, one-way ANOVA with Tukey’s multiple comparisons test, *p < 0.05, **p < 0.01,***p < 0.001; error bars ± SEM). The day 9 CD31^+^ endocardial starting population is used as the control (“D9 E 31^+^”). For all RT-qPCR analyses, expression values were normalized to the housekeeping gene *TBP*. Error bars represent SEM. *D*, day; *EndoMT*, endothelial-to-mesenchymal transition; *MACS*, magnetic-activated cell sorting; *E*, endocardium; *F*, bFGF; *B*, BMP4; *CH*, CHIR; *SB*, SB-431542.

To determine if the four-day CHIR/SB treatment altered the signalling requirements for EndoMT and VIC specification, we next manipulated BMP4/FGF and TGFβ2 signaling during this developmental step. For these experiments, the CHIR/SB treated cells were cultured for eight days with or without TGFβ2 in the presence or absence of BMP4 and bFGF (**Supplementary Fig 2a**). The resulting populations contained varying proportions of PDGFRβ^+^ cells and few, if any, CD31^+^ cells (**Supplementary Fig 2b**). The highest numbers of PDGFRβ^+^ cells were detected in cultures containing BMP4 and bFGF (**Supplementary Fig 2c**). The addition of TGFβ2 had no effect on the number of PDGFRβ^+^ cells generated. qRT-PCR analyses showed an upregulation of the expression of genes associated with the VIC lineage (*PRRX2*, *TIMP3, NR4A2)* in populations treated with BMP4 and bFGF (**Supplementary Fig 2d**). The levels of several of them (*PRRX2, TIMP3*) were further upregulated following treatment with TGFβ2. The four-day treatment with the combination of CHIR/SB/BMP4/bFGF did not alter the phenotype of the D9 VIC progenitor endocardial subpopulations, as the CD31^+^PDGFRβ**^+^**fraction generated under these conditions also gave rise to a higher frequency and total number of CD31^-^PDGFRβ^+^ mesenchymal derivatives than the CD31^+^PDGFRβ**^-^** subpopulation (**Supplementary Fig 3a–e**). Together, these findings show that a combination of BMP4, bFGF and TGFβ2 signalling functions to specify the VIC fate from CHIR/SB/BMP4/bFGF-treated CD31^+^PDGFRβ**^+^** endocardial cells.

### Heterogeneity of the hPSC-derived and fetal heart VIC populations

To further characterize the hPSC-derived VIC population, we analyzed it by scRNA-seq and compared the transcriptome of these cells to that of VIC populations isolated from the developing human heart. For the human heart populations, we micro-dissected the four valves from a normal fetal heart (gestational age, 105 days), enzymatically dispersed the tissue into single cells, and processed them for scRNA-seq. These analyses showed that, as in the mouse, the human valves contain *spongiosa* and *fibrosa* VIC subpopulations represented as distinct clusters in the UMAP plot (**Fig 5a).** Differential gene expression analysis revealed that both clusters expressed similar levels of *SOX9*, *CDH11*, and *VIM*. The *spongiosa* VIC cluster showed higher expression levels of *LUM, SERPINE2*, and APOE, whereas the *fibrosa* VIC cluster had higher levels of *PRRX2*, *TIMP3, COL1A1,* and *ITGA2* (*CD49b*) (**Fig 5b; Supplementary Table 1**).

**Figure 5.**
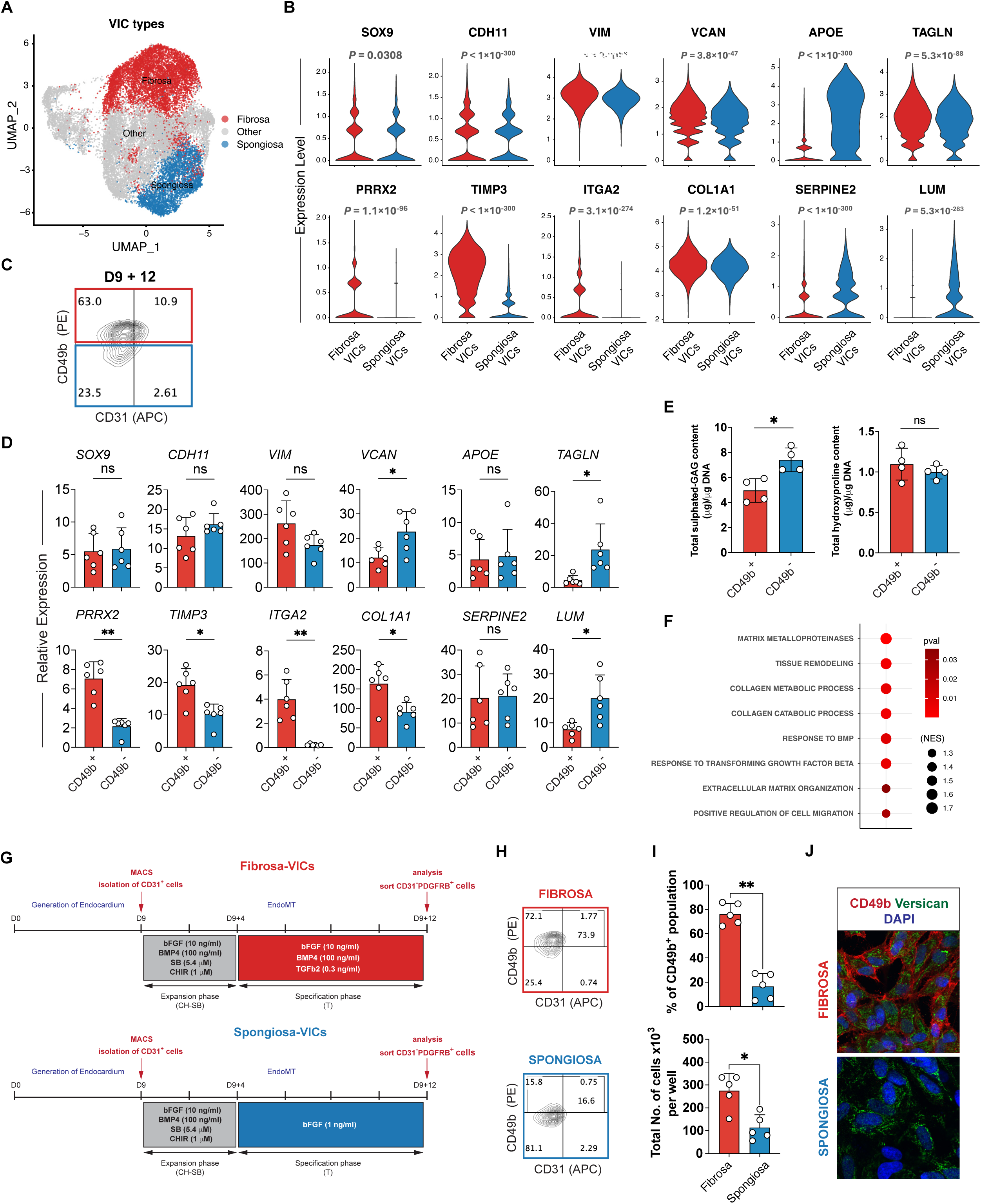
hPSC-derived VIC-like cells recapitulate the heterogeneity of primary human VICs. **(A)** UMAP plot of primary fetal valve populations showing clusters with enriched expression of *fibrosa* VICs genes (red) and *spongiosa* VIC genes (blue). **(B)** Violin plots showing expression of general mesenchymal markers (*SOX9*, *CDH11*, *VIM, COL1A1, VCAN, TAGLN*), *fibrosa* markers (*PRRX2*, *TIMP3*, *ITGA2*) and *spongiosa* markers (*LUM*, *SERPINE2*, *APOE*) in *fibrosa* and *spongiosa* clusters from (A). **(C)** Representative flow cytometric analysis of CD49 and CD 31 expression on a hPSC-derived VIC population generated with the protocol in **Supplementary Fig 3A**. **(D)** RT-qPCR analysis of the expression levels of general mesenchymal markers (*SOX9*, *CDH11*, *VIM, COL1A1, VCAN, TAGLN*), *fibrosa* markers (*PRRX2*, *TIMP3*, *ITGA2*) and *spongiosa* markers (*LUM*, *SERPINE2*, *APOE*) in the day 9+12 CD49b^+^ and CD49b^-^ cells generated with the protocol described in **Supplementary Fig 3A** and illustrated in (C) (n = 6, paired t-test *p < 0.05, **p < 0.01, ***p < 0.005). **(E)** Quantification of amount of glycosaminoglycans (GAG) and hydroxyproline secreted by CD49b^+^ and CD49^-^ cells from (C). The values are normalized to cell number represented by total amount of DNA from respective samples (n = 4, paired t-test *p < 0.05, **p < 0.01, ***p < 0.005). **(F)** GSEA showing pathways that are enriched in *fibrosa* compared to *spongiosa* VIC subpopulations from (A). **(G)** Schematics of the protocols used to generate and isolate hPSC-derived *fibrosa* and *spongiosa* VICs from CD31^+^ endocardial-like cells. **(H)** Representative flow cytometric analyses of *fibrosa* and *spongiosa* VIC populations generated with the protocols described in (G) showing the levels of CD49b expression in these populations. **(I)** Quantification of the frequency of CD49b^+^ cells in *fibrosa* and *spongiosa* VIC populations in (H)(n = 5, paired t-test *p < 0.05, **p < 0.01, ***p < 0.005). **(J)** Photomicrographs showing immunostaining of CD49b (red) and Versican (green) in *fibrosa* and *spongiosa* VIC populations generated with the protocols described in (G). The cells were co-stained with DAPI (blue) to visualize all cells and nuclei. Scale bars represent 50 μm. For all RT-qPCR analyses, expression values were normalized to the housekeeping gene *TBP*. Error bars represent SEM. *D*, day; *GAG*, glycosaminoglycan; *VIC*, valvular interstitial cells.

The higher expression of *ITGA2* in the *fibrosa* VIC cluster suggested that expression of the surface marker CD49b that it encodes may uniquely identify this subpopulation of VICs in the hPSC-derived D9+12 population. Flow cytometry analyses revealed that our D9+12 VIC-like population was indeed heterogeneous for expression of CD49b and contained both positive and negative cells (**Fig 5c; Supplementary Fig 4a**). To determine if CD49b expression distinguished the two VIC subtypes, we isolated the CD49b^+^ and CD49b^-^ fractions from the D9+12 population and analyzed them by qRT-PCR for molecular signatures of both the *spongiosa* and *fibrosa* VICs. These analyses showed that the CD49b^+^ fraction expressed higher levels of the *fibrosa* VIC markers *PRRX2*, *TIMP3, COL1A1* and *ITGA2,* whereas the CD49b^-^ fraction had higher levels of expression of the *spongiosa* VIC marker *LUM* (**Fig 5d**). These expression profiles suggest that CD49b can be used to distinguish and isolate *fibrosa*- and *spongiosa*-like VICs. To test the functional implications of these molecular differences, we performed biochemical analyses to measure the quantity of hydroxyproline, a major component of collagen, and glycosaminoglycans (GAGs) produced by CD49b^+^ and CD49b^-^ VIC-like cells. While both populations produced similar quantities of collagens (**Fig 5e**), the CD49b^-^ VIC-like cells generated more GAGs, supporting the notion that these cells bear more phenotypical resemblance to *spongiosa* VICs.

The protocol depicted in **Supplementary Fig 3a** generated populations that consistently contained a higher proportion of CD49b^+^ than CD49b^-^ cells (**Fig 5c**), suggesting that the pathway agonists used skewed differentiation towards a *fibrosa* VIC-like fate. Given this, we next investigated whether the protocol could be modified to give rise to a higher proportion of GAG-producing *spongiosa* VIC-like cells. Using GSEA, we first sought to identify signalling pathways that were enriched in *fibrosa* VICs compared to *spongiosa* VICs in the human fetal heart valve scRNA-seq dataset. These analyses showed that pathways related to “response to BMP” and “response to transforming growth factor beta” were enriched in *fibrosa* VICs compared with *spongiosa* VICs (**Fig 5f**). Since our VIC protocol contains both BMP4 and TGFβ2, we removed these factors and cultured the cells in FGF alone in an attempt to enhance the proportion of CD49b^-^ (*spongiosa*) VIC-like cells generated (**Fig 5g**). Flow cytometric analyses showed that the population generated in the presence of only FGF contained a significantly higher proportion of CD49b^-^ cells than the one cultured with TGFβ2 and BMP4 in addition to FGF (**Fig 5h and i; Supplementary Fig 4b**). Immunostaining analyses were consistent with the findings and showed that only the cells treated with FGF, TGFβ2 and BMP4 expressed CD49b (**Fig 5j**).

### Transcriptomic analysis of hPSC-derived VICs

To characterize in detail the molecular profiles of the hPSC-derived VIC-like cells obtained from the two protocols depicted in **Fig 5g**, we carried out scRNA-seq analyses. The cells obtained from the *fibrosa* and *spongiosa* protocols were labelled with a cell multiplexing method (10x Genomics 3’CellPlex kit with Feature Barcode technology)^23^ before sequencing. As shown in **Fig 6a**, the cells derived from the two protocols were found to cluster separately on UMAP projection. Differential gene expression analysis between the two clusters showed that *fibrosa* VIC-like cells express higher levels of *PRRX2* and *ITGA2*, whereas *spongiosa* VIC-like cells express higher levels of *LUM*, *TAGLN*, *VCAN* and *APOE* (**Fig 6b; Supplementary Table 2**). These expression profiles are consistent with the interpretation that the *fibrosa* and *spongiosa* protocols yield distinct subtypes of VIC-like cells.

**Figure 6.**
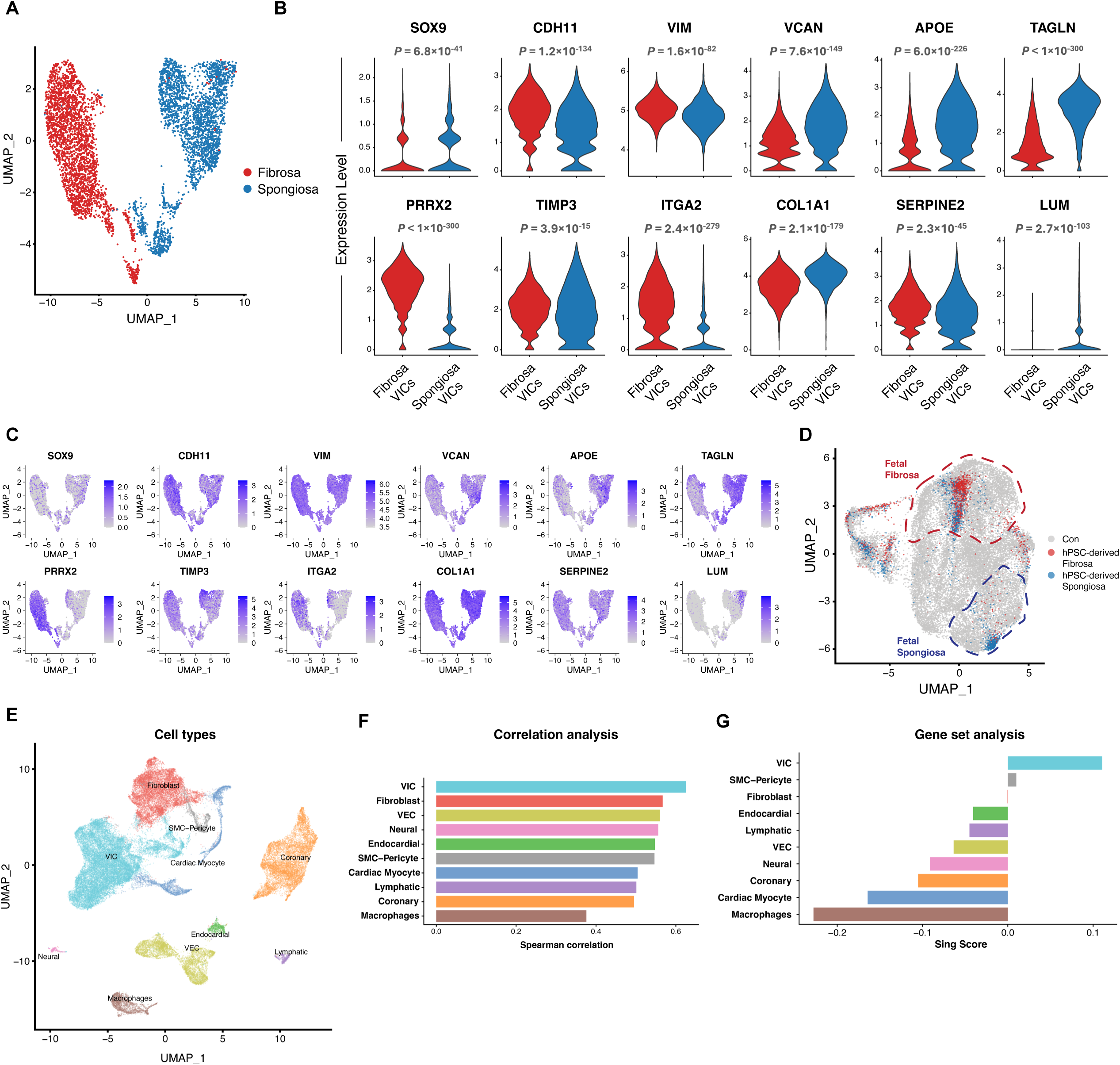
scRNA-Seq analysis of hPSC-derived VICs. **(A)** UMAP plot of hPSC-derived populations of putative *fibrosa* VICs (red) and *spongiosa* VICs (blue) generated with the protocols in Figure 5G and labeled with cell multiplexing oligos prior to sequencing. **(B)** Violin plots showing expression of general mesenchymal markers (*SOX9*, *CDH11*, *VIM, COL1A1, VCAN, TAGLN*), *fibrosa* markers (*PRRX2*, *TIMP3*, *ITGA2*) and *spongiosa* markers (*LUM*, *SERPINE2*, *APOE*) in *fibrosa* and *spongiosa* clusters from (A). **(C)** UMAP plots showing expression of general mesenchymal markers (*SOX9*, *CDH11*, *VIM, COL1A1, VCAN, TAGLN*), *fibrosa* markers (*PRRX2*, *TIMP3*, *ITGA2*) and *spongiosa* markers (*LUM*, *SERPINE2*, *APOE*) in *fibrosa* and *spongiosa* clusters from (A). **(D)** Reference mapping analysis of hPSC-derived *fibrosa* (red) and *spongiosa* (blue) VICs projected on the primary human fetal UMAP with primary *fibrosa* and *spongiosa* VIC clusters outlined with red and blue dashed lines, respectively. **(E)** UMAP plots of fetal heart populations colored by cluster identity. **(F)** Correlation analysis of hPSC-derived VICs (combined *fibrosa* and *spongiosa*) versus fetal heart populations. **(G)** Gene set scoring analysis of hPSC-derived VICs. *hPSC*, human pluripotent stem cells; *UMAP*, day; *VIC*, valvular interstitial cells.

While the populations are distinct, UMAP plots revealed that a small proportion of cells generated from the *spongiosa* protocol expressed the *fibrosa* VIC markers *PRRX2*, *TIMP3*, and *ITGA2*, suggesting a degree of overlap (**Fig 6c**). Comparison of the hPSC-derived populations to those from the fetal heart showed that the *in vitro*-derived *fibrosa* and *spongiosa* VIC-like cells largely aligned molecularly with the respective subtypes from the fetal tissue. There were, however, some hPSC-derived cells that clustered with primary human VICs of the opposite subtype, suggesting that not all the *in vitro*-derived VICs are properly specified (**Fig 6d**). Additionally, a proportion of hPSC-derived VIC-like cells obtained from both protocols were characterized by an intermediate expression profile with molecular features of both *fibrosa* and *spongiosa* VICs (**Fig 6d**).

To further assess the transcriptomic fidelity of hPSC-derived VIC-like cells, we compared their expression profile to that of various cardiac cell populations obtained from the primary human fetal tissue. Cell type assignment using clustering and canonical markers revealed that most cell types expected to be present in the fetal heart were represented, including cardiomyocytes, endocardial cells, coronary vascular cells, VICs, VECs, fibroblasts, smooth muscle cells/pericytes, neural cells, lymphatic cells, and macrophages (**Fig 6e**). Spearman’s correlation analysis (see Methods) showed that hPSC-derived VIC-like cells share a global gene expression profile similar to their counterparts in the primary human fetal heart (**Fig 6f**). Notably, the hPSC-derived VIC-like cells were transcriptionally more similar to primary human VICs than to any other cell type identified in the primary human fetal heart. To further evaluate similarities between hPSC-derived VIC-like cells and primary human fetal VICs, we performed GSEA to calculate the cell type-specific signature enrichment in hPSC-derived VICs. Briefly, we first established a gene expression signature for each subpopulation in the human fetal heart by identifying the top 100 significantly up- or down-regulated genes. We then used the SingScore method to assess the transcriptomic similarity of hPSC-derived VICs to each of these signatures ^24^ (see Methods for details). This analysis confirmed the high degree of transcriptomic similarity between hPSC-derived VICs and fetal VICs, in contrast to all the other fetal cardiac populations (**Fig 6g; Supplementary Fig 5**). Taken together, these scRNA-seq analysis findings demonstrate that hPSC-derived VIC-like cells have molecular profiles similar to those of primary human fetal VICs.

### hPSC-derived VICs produce human tissue-engineered extracellular matrices

*In situ* tissue engineering has the potential to overcome the limitations of mechanical and xenogeneic heart valve replacements currently used in clinical practice ^7^. A hybrid material of human cells that are seeded on a polymeric scaffold to produce human ECM has demonstrated a reduced incidence of thrombogenic and immunogenic adverse events in pre-clinical models ^25–27^. To serve as a potential human cell source for *in situ* tissue-engineered heart valve manufacturing, hPSC-derived VICs must be able to secrete, remodel, and mature the ECM sufficiently. To test their ECM secretion capacity, hPSC-derived *fibrosa and spongiosa* VICs were seeded onto polyglycolic acid/poly-4-hydroxybutyrate polymeric scaffolds and cultured for 4 to 6 weeks in the presence of 0.3 ng/mL or 1.0 ng/mL TGFβ2 to stimulate ECM production. In preliminary experiments in which TGFβ2 was omitted, no appreciable ECM formation was observed, and the scaffolds degraded over the 4-week culture period (n = 2, data not shown), confirming the necessity of TGFβ2 supplementation for maintaining scaffold integrity and promoting matrix deposition. These human tissue-engineered matrices (hTEMs) were then decellularized and subjected to histological, immunohistochemical and biochemical analyses (**Fig 7a**). After 4 weeks of dynamic culture, all the hTEMs displayed neo-tissue deposition and homogeneous matrix distribution (**Fig 7b and c**). Hematoxylin and eosin staining demonstrated more abundant tissue deposition and a more compact and organized neo-tissue structure in the *spongiosa* VIC group, compared with the *fibrosa* VIC group (**Fig 7c**). Immunohistochemistry confirmed the presence of a collagen-rich matrix comprised of collagen types 1 and 3 in all tissues (**Fig 7c**). Tissues generated from *spongiosa* VICs exhibited a thicker collagen layer than those produced from *fibrosa* VICs. The presence of these collagens is important as they are the most abundant ECM proteins found in native heart valves. However, evaluation of picrosirius red-stained samples under polarizing light revealed that the collagen fibers remained disorganized, indicative of incomplete collagen maturation (**Supplementary Fig 6**).

**Figure 7.**
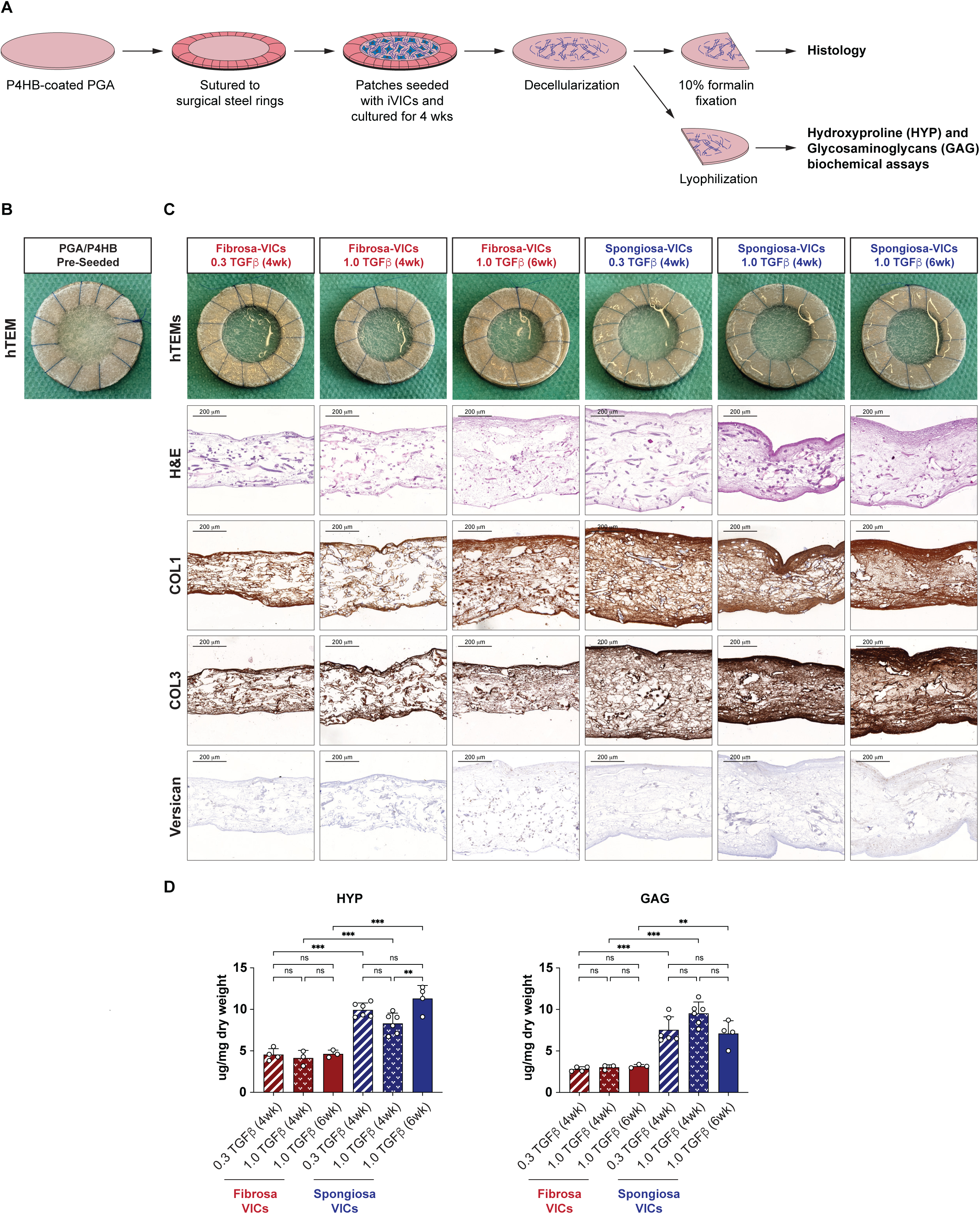
hPSC-derived VICs produce human tissue-engineered extracellular matrices. **(A)** Schematic depiction of the hTEM production process and analysis of hPSC-derived VIC ECM secretion. **(B)** Representative photomicrograph of the polyglycolic acid/poly-4-hydroxybutyrate scaffold sutured onto a stainless metal ring before cell seeding and *in vitro* culture. **(C)** Top panel: representative photomicrographs of polyglycolic acid/poly-4-hydroxybutyrate patches seeded with hPSC-derived *fibrosa* and *spongiosa* VICs, cultured for four weeks in the presence of 0.3 or 1 ng/ml of TGFβ2. Bottom panels: representative qualitative images of hTEMs subjected to histology staining using H&E and immunohistochemistry of collagen 1, collagen 3, and versican. Scale bars indicate 200 μm. **(D)** Hydroxyproline (HYP) and glycosaminoglycan (GAG) quantification using biochemical assays. (one-way ANOVA with Tukey’s multiple comparisons test, *p < 0.05, **p < 0.01,***p < 0.001; error bars ± SEM). *COL1*, collagen 1; *COL3*, collagen 3; *GAG*, glycosaminoglycan; *H&E*, hematoxylin and eosin; *hTEM*, human tissue-engineered matrix; *HYP*, hydroxyproline; *VIC*, valvular interstitial cell.

Similar to collagen, versican staining was most visible in hTEMs seeded with *spongiosa* VICs. Most of the versican deposition was found in the areas of dense collagen matrix (**Fig 7c**). Quantification of hydroxyproline, which measures collagen content in tissues, and glycosaminoglycans via biochemical analyses confirmed the histological findings and demonstrated higher collagen and glycosaminoglycan content in the *spongiosa* tissue compared to the *fibrosa* tissue (**Fig 7d**). Interestingly, the concentration of TGFβ2 (0.3 ng/mL versus 1.0 ng/mL) did not significantly impact the quantity of collagen or glycosaminoglycan secreted in either group (**Fig 7d**). However, on histological appearance, higher concentrations of TGFβ2 were associated with denser ECM deposition in the *spongiosa* VIC group (**Fig 7c**). Overall, hTEMs produced with hPSC-derived *spongiosa* VICs and treated with 1.0 ng/mL TGFβ2 showed the most pronounced ECM stratification. Taken together, these findings demonstrate that hPSC-derived VICs can generate collagen-containing tissues and show that those directed towards a *spongiosa* lineage exhibit a superior ability to synthesize ECM compared to those differentiated into *fibrosa* cells.

### hPSC-derived VICs recapitulate the pathogenesis of Noonan syndrome valve disease

Next, we sought to determine if the hPSC-derived VIC populations could be utilized to model valve disease. Specifically, we chose to model Noonan syndrome, a developmental disorder that arises from gain-of-function variants in the RAS-MAPK signalling pathway. Patients with Noonan syndrome frequently develop congenital heart defects, most commonly in the form of pulmonary valve stenosis. Past attempts to model this pathology *in vivo* have shown that mice exhibit enlarged endocardial cushions, have dysregulated EndoMT, and that the resulting mesenchymal cells have hyperactivation of phospho-ERK1/2 ^28, 29^. To determine if our hPSC-derived VICs could model valve disease in Noonan syndrome and recapitulate known aspects of its pathology, we utilized CRISPR-Cas9 to edit a non-mutant induced pluripotent stem cell (iPSC) line and introduced the pathogenic *PTPN11^N308D/N308D^* variant (**Fig 8a**). This *PTPN11* N308D variant is the most common Noonan syndrome allele and is strongly associated with pulmonary valve stenosis ^30, 31^.

**Figure 8.**
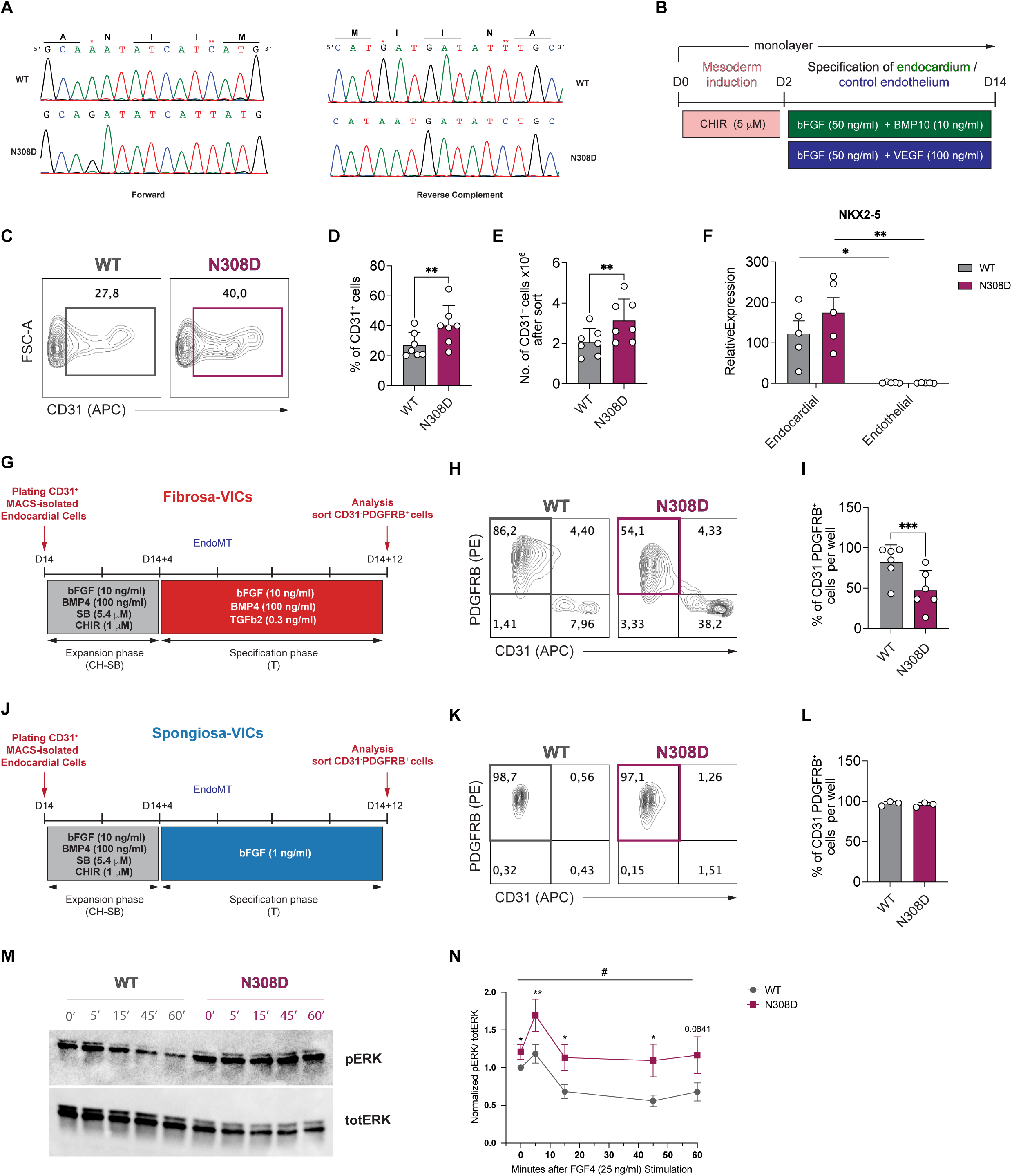
hPSC-derived VICs recapitulate the pathogenesis of Noonan syndrome valve disease. **(A)** Sanger sequences of WT and *PTPN11^N308D/N308D^* iPSC lines (*denotes N308D mutation site, **silent PAM edit). **(B)** Schematic of the protocol used for the generation and isolation of iPSC-derived endocardial-like cells by sequential induction with CHIR, bFGF and BMP10, as well as control endothelial cells by induction with CHIR, bFGF and VEGF. The cells were cultured in a monolayer format throughout. **(C)** Representative flow cytometric analyses of CD31 expression on endocardial cells (day 14) generated from the WT and *PTPN11^N308D/N308D^* iPSC lines with the protocol in (B). **(D)** Quantification of the frequency of CD31^+^ cells generated from WT and *PTPN11^N308D/N308D^* mutant iPSC-derived cells at day 14 of culture. **(E)** Quantification of the total number of CD31^+^ cells isolated from WT and *PTPN11^N308D/N308D^*mutant iPSC-derived endocardial cultures at day 14 by CD31 magnetic bead sort (MACS) (n = 7, paired t-test *p < 0.05, **p < 0.01, ***p < 0.005). **(F)** RT-qPCR analysis of the expression levels of *NKX2-5* in the day 14 CD31^+^ endocardial or endothelial cells generated from the WT and N308D hPSCs (n = 5, two-way ANOVA with multiple comparison analysis using Sidak’s post hoc test *p < 0.05, **p < 0.01, ***p < 0.005). **(G)** Schematic of the protocol used to generate and isolate iPSC-derived *fibrosa* VICs from day 14 CD31^+^ endocardial-like cells. **(H)** Representative flow cytometric analyses of CD31 and PDGFRβ expression on VICs generated from WT and *PTPN11^N308D/N308D^* iPSCs following 12 days of culture (D14+12) using the protocol described in (G). **(I)** Quantification of the frequency of CD31^-^PDGFRβ^+^ cells generated from the WT and *PTPN11^N308D/N308D^* iPSC lines following 12 days of culture (D14+12) using the protocol described in (G) (n = 7, paired t-test *p < 0.05, **p < 0.01, ***p < 0.005). **(J)** Schematic of the protocol used to generate and isolate iPSC-derived *spongiosa* VICs from day 14 CD31^+^ endocardial-like cells. **(K)** Representative flow cytometric analyses of of CD31 and PDGFRβ expression on VICs generated from WT and *PTPN11^N308D/N308D^*iPSCs following 12 days of culture (D14+12) using the protocol described in (J). **(L)** Quantification of the frequency of CD31^-^PDGFRβ^+^ cells generated from the WT and *PTPN11^N308D/N308D^* iPSC lines following 12 days of culture (D14+12) using the protocol described in (J) (n = 3, paired t-test *p < 0.05, **p < 0.01, ***p < 0.005). **(M)** Western blot analysis of ERK1/2 phosphorylation in WT and N308D *fibrosa* VICs following stimulation with FGF4 for 5, 15, 45 and 60 minutes. **(N)** Quantification of ERK1/2 phosphorylation in (M) (n = 6, paired t-test *p < 0.05, **p < 0.01; # denotes an overall ANOVA p-value < 0.05; error bars ± SEM.

For these studies, our previously published embryoid body-based endocardial protocol was modified to a feeder-free, monolayer culture system ^32^. With this approach, iPSCs were grown without feeders on Matrigel and mesoderm was induced through activation of WNT signalling (CHIR). This mesoderm was specified to an endocardial fate with BMP10 and bFGF (**Fig 8b**). The sorted CD31^+^ endocardial population was further directed to undergo EndoMT towards either *fibrosa* or *spongiosa* VICs using the approach described above (**Fig 8g and j**). Applying this methodology to non-mutant and N308D iPSC lines, we found that the *PTPN11* N308D line reproducibly generated a higher frequency and total number of CD31^+^ cells than the non-mutant cells (**Fig 8c–e**). To validate the endocardial identity of the CD31^+^ cells thus generated and isolated, we examined their expression of the cardiac transcription factor NKX2-5. RT-qPCR analyses revealed that for both the WT and *PTPN11* N308D lines, BMP10-derived endocardial cells upregulated expression of *NKX2-5* compared to the control endothelial population generated with VEGF (**Fig 8f**).

Next, we induced EndoMT in these endocardial populations, directing them towards either *fibrosa*-like or *spongiosa*-like VICs (**Fig 8g and j**). After 12 days of EndoMT, we observed a marked decrease in EndoMT efficiency for the *fibrosa-*like VIC protocol (**Fig 8h and i**), but not the *spongiosa-*like VIC protocol (**Fig 8k and l**). Since the *fibrosa* protocol exhibited reduced EndoMT efficiency, we focused on this population to model Noonan syndrome. Specifically, we asked if these *fibrosa-*like VICs could recapitulate one of the molecular hallmarks of Noonan syndrome, namely increased RAS/MAPK signalling. To test this, we first isolated D14+12 *fibrosa-*like VICs via negative selection and serum-starved the cells overnight (see Methods for details). We then stimulated these VICs with 25 ng/mL of FGF4 — a growth factor known to be involved in proper valve mesenchyme development ^33, 34^ — for 5, 15, 45, and 60 minutes. Following stimulation, western blot analysis was performed on the cell lysates and probed for phosphorylated ERK1/2 (pERK) (**Fig 8m**). These analyses revealed that pERK signalling was significantly higher in the *PTPN11* N308D VICs than non-mutants at all time points, including at baseline (**Fig 8n**). In particular, we found that N308D VICs displayed an increased magnitude of pERK activation following FGF4 stimulation at each time point. The non-mutant VICs showed an initial increase in pERK at 5 min, followed by a rapid negative regulation by 15 min, reducing pERK levels significantly below its unstimulated baseline. This trend continued through the 60-minute timeframe. The *PTPN11* N308D VICs similarly downregulated pERK within 15 minutes of FGF4 stimulation. However, in contrast to the non-mutant cells, the pERK levels in the mutant cells declined only to those of the unstimulated cells and not below (**Fig 8m and n**). Collectively, these data suggest that the N308D-hPSCs exhibit abnormal valvulogenesis at multiple stages, including increased endocardial cell specification and diminished EndoMT potential. Furthermore, the mutant *fibrosa-*like VICs recapitulate a primary molecular defect of Noonan syndrome—increased pERK activity and magnitude of response following FGF4 stimulation.

## Discussion

The ability to generate functional human VICs from hPSCs would provide an unlimited source of these cells for engineering biological valves for valve replacement therapy and for modelling diseases that impact valve function. Several previous studies have reported on the generation of putative VIC-like cells from hPSCs ^35, 36^. In the first of these, Neri *et al*. demonstrated that a heterogenous endothelial cell population specified from hPSCs could undergo EndoMT and generate mesenchymal cells ^35^. However, as the identity of the population was not firmly established, it is unclear what proportion represents VIC lineage cells. In a very recent study, Cai *et al*. described a 6-day differentiation protocol that yielded mesenchymal cells that showed transcription profiles suggestive of VIC development ^36^. While these molecular analyses are consistent with the generation of VIC-like cells, their rapid specification in culture (6 days) and the lack of demonstration of an endocardial intermediate raises questions as to their origin and their relationship with valve-like cells that emerge through the normal developmental trajectory. In the present study, we used a developmental biology-guided approach to establish a protocol that promotes the generation of mesenchymal cells from hPSC-derived endocardial cells that display the following defining characteristics of the *bona fide* VICs. First, we show that the hPSC VIC lineage derives from a subpopulation of endocardial cells, defined by the co-expression of CD31 and PDGFRβ, that displays properties of cushion endocardium. Second, our findings identify distinct subpopulations of hPSC-derived *fibrosa*- and *spongiosa*-like VICS, define key signalling pathways that promote their development and characterize a surface marker to distinguish between them. Third, we demonstrate that the two VIC subpopulations display molecular profiles similar to their counterparts isolated from primary human fetal valve tissue.

Studies in different model organisms have shown that the ability of endocardial cells to undergo EndoMT is tightly restricted to the endocardium of the atrioventricular and outflow tract cushion regions, suggesting molecular and functional differences between chamber and cushion endocardium^21, 37^. In this study, we recapitulated some of these differences in the hPSC-derived endocardial population and identified PDGFRβ as a potential marker of the *in vitro* equivalent of cushion endocardium. This lineage assignment is based on the observation that the PDGFRβ^+^CD31^+^ endocardial subpopulation displayed characteristic features of cushion endocardium, including elevated expression of genes associated with VIC progenitors and the capacity to undergo EndoMT and generate mesenchymal progeny with VIC-like properties. Additionally, these cells showed a calcium response to exogenous ATP, a property recently reported to distinguish atrioventricular canal cushion endocardium from chamber endocardium in the zebrafish ^14^. In contrast, the PDGFRβ^-^CD31^+^ cells expressed higher levels of NRG1, a marker associated with chamber endocardium and showed little, if any, EndoMT potential or calcium response to ATP. Collectively, these observations demonstrate that the hPSC-derived cushion endocardial cells are present in the endocardial population by D9 of differentiation and as such raise interesting questions regarding the stage at which this fate is established and the signalling pathways regulating this developmental decision. Our observation that BMP10 signalling is required for the generation of the hPSC endocardial population ^9^ demonstrates that this pathway plays some role in the specification of the PDGFRβ^+^CD31^+^ cells. Whether or not other pathways regulate the specification of the PDGFRβ^+^CD31^+^ vs the PDGFRβ^-^CD31^+^ subpopulations remains to be determined.

A critical step in valve formation is the generation of VICs from the cushion endocardium through a process known as EndoMT. Previous studies have shown that EndoMT is mediated, in part, by BMP and TGFβ signalling ^9, 38, 39^. However, under these conditions, the process is inefficient, yielding mixed populations containing 25-30% of mesenchymal cells on average. This low yield may indicate that the D9 PDGFRβ^+^CD31^+^ endocardial cells are not optimally primed to undergo EndoMT or that other pathways are involved in this process. Additional insights into the regulation of this process have come from studies in model organisms that showed that Wnt signalling plays a role in cushion formation, EndoMT, and VIC formation in both mice and zebrafish^22, 40^. Our findings show that by activating the Wnt signalling pathway while simultaneously inhibiting the TGF beta pathway in PDGFRβ^+^CD31^+^ endocardial cells, we markedly enhance the efficiency of EndoMT. This enhancement is evident in both the purity of the generated cell population and the overall output of VIC-like cells. These findings underscore the parallel regulatory mechanisms governing valve development in humans and model organisms, shedding light on the intricate processes involved in this crucial stage of development. With this advance, it is possible to easily generate tens of millions of VIC-like cells that can be cryopreserved for different applications.

The outcome of our molecular profiling showing a high level of transcriptional fidelity between the hPSC-derived and primary human VICs provides strong evidence that we have generated *bona fide* VICs rather than generic mesenchymal cells. These analyses also revealed that the fetal human VIC population contains distinct collagen- and glycosaminoglycan-expressing interstitial cell clusters, which we propose correspond to the *fibrosa* and *spongiosa* VICs described in the mouse ^4^. The heterogeneity identified here likely reflects the subpopulations of VICs that reside within different layers of the heart valve leaflet, each characterized by unique molecular signatures and specific protein secretion patterns. Segregation of VIC subsets within the valve leaflet contributes to the stratified nature of the valve’s ECM, a feature essential to its function and durability ^4, 41^. Given this, access to distinct hPSC-VIC populations is important for achieving our ultimate goal of engineering a biological living valve. To address the challenge of generating VIC subsets from hPSC-derived endocardial cells, we manipulated BMP and TGFβ signalling in the D9+4 population as these pathways were identified as potential drivers of the *fibrosa* fate *in vivo*. With this approach, we successfully specified distinct VIC populations that expressed molecular similarities to the *fibrosa* and *spongiosa* VICs identified in the fetal VICs. In addition to the molecular differences, these subpopulations displayed divergent matrix secretion profiles. When cultured on tissue culture plastic, both populations produced similar levels of collagen, but CD49b⁻ (*spongiosa*-like) cells secreted significantly more GAGs. Although we did not detect a significant difference in hydroxyproline content between *fibrosa-like* and *spongiosa-like* cells, this result is consistent with native valve composition. Indeed, while the *fibrosa* is traditionally considered collagen-rich, the *spongiosa* also contains appreciable amounts of collagen—particularly during development and in the neonatal period ^42^. Thus, comparable collagen production by *spongiosa-like* cells is biologically plausible. In contrast, GAGs are largely confined to the *spongiosa*, making the observed GAG enrichment in CD49b⁻ cells more reflective of layer-specific matrix specialization. These findings align with prior histological studies demonstrating collagen presence in both *fibrosa* and *spongiosa* layers, with GAGs primarily localized to the *spongiosa* ^42, 43^.

When seeded in tissue-engineered scaffolds, *spongiosa* VICs secreted more ECM overall than the *fibrosa* VICs. Based on native valve composition, we anticipated that *spongiosa* VICs would be enriched in GAGs and *fibrosa* VICs in collagen. Instead, the *spongiosa* VICs produced significantly more GAGs and more collagen. The higher glycosaminoglycan content in the *spongiosa* tissues may reflect the fact that these cells are not mature and likely represent an early fetal stage of development, a stage at which the glycosaminoglycans are the dominant ECM produced ^35^. Picrosirius red staining with polarized light microscopy further revealed that although abundant collagen was present in the *spongiosa* VIC group, the absence of characteristic green or orange birefringence indicated that the fibers remained immature and lacked the organized architecture of mature collagen. Overall, these findings suggest that while the ECM produced by *spongiosa* VICs does not yet display the structural hallmarks of maturity, these cells’ abundant output of both GAGs and collagen in engineered scaffolds represents an important precursor to ECM maturation and a key step towards functional tissue development.

Because these findings rely on the integrity of the underlying scaffold, it is important to note that our detergent-based decellularization protocol effectively removes cellular components while preserving ECM structure ^44^, though it may deplete key ECM components such as proteoglycans and GAGs, as evidenced by the absence of versican in our samples ^45^. Additionally, while structural integrity is maintained, biochemical composition and mechanical properties may be altered, potentially impacting functionality *in vivo*. Despite these drawbacks, this method remains widely used in clinical scaffold preparation to prevent immune rejection and support cell infiltration and tissue remodeling.

To further demonstrate the utility of hPSC-derived VICs, we sought to recapitulate known aspects of valve disease in Noonan syndrome. Transgenic mouse studies have shown that Noonan syndrome gain-of-function mutations in the *PTPN11* gene — which encodes the protein tyrosine phosphatase SHP2 — result in enlarged endocardial cushions ^28^. In humans, *PTPN11* mutations account for ∼50% of Noonan syndrome cases and are associated with high rates of pulmonary valve stenosis ^46^. Our demonstration that Noonan syndrome hiPSCs have increased differentiation towards the endocardial lineage is consistent with the *in vivo* findings of enlarged endocardial cushions. Interestingly, we also show that Noonan syndrome endocardial cells demonstrate defective EndoMT towards *fibrosa* VICs, suggesting there may be subtype-specific contributions to Noonan syndrome valve stenosis during development. Beyond the differences in differentiation efficiency, our observation that the Noonan syndrome *fibrosa* VICs exhibited increased phosphorylation of ERK1/2, and aberrant negative regulation of ERK compared with non-mutant cells is consistent with studies in the mouse showing enhanced ERK1/2 signalling in Noonan syndrome endocardial cushion mesenchymal cells ^28, 29, 47^.

Sex and age are important considerations in the interpretation of findings from this study. All hPSC-derived VICs were generated from male hPSC lines, and sex-specific differences in VIC phenotype or function may not be fully captured. Furthermore, our hPSC-derived VICs likely resemble a fetal or early postnatal phenotype; further studies will be needed to determine how they compare to VICs from aged human valves, particularly with respect to matrix remodeling and calcification.

In conclusion, this study has successfully identified a distinct CD31^+^PDGFRβ^+^ hPSC-derived endocardial population that displays properties of cushion endocardium, including the ability to undergo EndoMT and give rise to VIC-like cells. Additionally, it has demonstrated that, as observed in murine and human fetal heart valve tissues, the hPSC-derived VIC populations contain distinct *fibrosa* and *spongiosa* subsets. Together, these advances bring us an important step closer to creating *in vitro* models for valvulogenesis and heart valve diseases and to developing engineered heart valve tissues that more faithfully mirror the intricate cellular makeup of human heart valves.

## Supplemental Methods

### Generation and maintenance of HES3-NKX2-5eGFP/w and HES2:WT hESC lines

HES3-NKX2-5^eGFP/w^ (karyotype: 46, XX) reporter hPSCs^48^ and HES2:WT hPSCs ^49^ (karyotype: 46, XX) were maintained on irradiated mouse embryonic fibroblasts in hPSC culture media consisting of DMEM/F12 (Cellgro), penicillin/streptomycin (1%, ThermoFisher), L-glutamine (2 mM, ThermoFisher), non-essential amino acids (1x, ThermoFisher), β-Mercaptoethanol (55 μM, ThermoFisher) and KnockOut^TM^ serum replacement (20%, ThermoFisher) at 37 °C in a normoxic environment (5% CO_2_)as described previously ^50^.

### Generation and maintenance of WT and Noonan syndrome iPSCs

Wild-type human iPSC line WTC11 was obtained from the Coriell Institute (XY, mono-allelic ACTN2-mEGFP, #AICS-0075-085). A homozygous edit of *PTPN11* N308D/N308D via CRISPR was designed and executed by Synthego. These iPSC lines were maintained in 6-well plates coated with a thin layer of Matrigel (6.6% v/v, Corning), diluted in DMEM (ThermoFSisher) and supplemented with penicillin/streptomycin (5%, ThermoFisher). iPSCs were maintained and fed daily with iPSC medium, consisting of mTesR Plus (Stem Cell Technologies) supplemented with penicillin/streptomycin (1%, ThermoFisher). Passaging of iPSCs was performed every 3 to 4 days when cells reached ∼80–90% confluency by first dissociating with Accutase (Innovative Cell Technologies, Inc) and then replating at a 1:10 ratio with iPSC medium, supplemented with Thiazovivin (2 µM, Selleckchem) for the first 24 hours. iPSCs were incubated at 37 °C in a normoxic environment (5% CO_2_).

### Generation of endocardial/endothelial cells from HES3-NKX2-5eGFP/w and HES2:WT hESC lines

To generate endocardial/endothelial cells, hPSCs were differentiated to cardiogenic mesoderm induced with 5 ng/ml BMP4 and 3 ng/ml activin A using the protocol described in our previous work ^9^. To initiate differentiation, hPSCs at 80%-90% confluence were dissociated into single cells (TrypLE, ThermoFisher) and re-aggregated to form embryonic bodies (EBs) at a cell density 5×10^5^ cells/ml in StemPro-34 media (ThermoFisher) supplemented with penicillin/streptomycin (1%, ThermoFisher), L-glutamine (2 mM, ThermoFisher), ascorbic acid (50 μg/ml, Sigma), monothioglycerol (50 μg/ml, Sigma), transferrin (150 μg/ml, ROCHE), ROCK inhibitor Y-27632 (10 μM, TOCRIS) and rhBMP4 (1 ng/ml, R&D). For EB generation, the cultures were rotated for 18 h on an orbital shaker (MaxQ 2000 shaker, Thermofisher) in 6-cm petri dishes (VWR) at 60 RPM. Following the rotation step, the EBs were transferred to fresh 6-cm petri dishes (VWR) in mesoderm induction media consisting of StemPro-34 supplemented with penicillin/streptomycin (1%), L-glutamine (2 mM), ascorbic acid (50 μg/ml), monothioglycerol (50 μg/ml), transferrin (150 μg/ml), rhBMP4 (5 ng/ml, R&D), rhActivinA (3 ng/ml, R&D) and rhbFGF (5 ng/ml, R&D). At day 3, the EBs were dissociated with TrypLE for 3 min at 37 °C and the cells plated in 6-well flat bottom plates (Corning) coated with Matrigel (25% v/v, Corning) at a density of 10^6^ cells per well. The monolayers were cultured in StemPro-34 supplemented with penicillin/streptomycin (1%, ThermoFisher), L-glutamine (2 mM, ThermoFisher), ascorbic acid (50 μg/ml, Sigma), monothioglycerol (50 μg/ml, Sigma), transferrin (150 μg/ml, ROCHE) and rhbFGF (50 ng/ml, R&D). For endocardial induction, rhBMP10 (10 ng/ml) was added to the media from day 5 to day 9. For generation of control endothelial cells, VEGF (100 ng/mL, R&D) was added to the differentiation media from day 3 to day 9. The cultures were incubated in a low oxygen environment (5% CO_2_, 5%O_2_, 90% N_2_) and the media was changed every two days. At day 9, HES3-NKX2-5^eGFP/w^-derived endocardial/endothelial cells were analyzed and isolated based on the expression of NKX2-5:GFP and CD31. The endocardial cells generated from the non-transgenic HES2-WT hPSC line were analyzed and isolated as CD31^+^ populations. For dissociation, the monolayers were incubated in collagenase type 2 (1 mg/ml, Worthington) in HANK’s buffer (Gibco) at 37 °C for one hour.

### Generation of mesenchymal / VIC-like cells from hESC-derived endocardial cells

Day 9 endocardial bulk cultures (HES3-NKX2-5^eGFP/w^ or HES2:WT lines) were dissociated with collagenase type 2 as described above. Single-cell suspensions were enriched for CD31+/CD34+ cells by magnetically activated cell sorting (MACS, Miltenyi, 130-146-702) to a purity of over 95%. Cells were incubated with anti-CD31 or anti-CD34 microbeads for 30 minutes at 4 °C in base media supplemented with DNAse (1U/ml, Millipore)(10μl microbeads/5×10^6^ cells in 100 μl of media) and purified by MS or LS columns. After the sort, CD31+/CD34+ endocardial cells were plated in a 12-well flat bottom microplate (Corning) coated with Matrigel (25% v/v, Corning) at a density 3×10^5^ cells per well and cultured for 8 or 12 days in StemPro-34 supplemented with penicillin/streptomycin (1%, ThermoFisher), L-glutamine (2 mM, ThermoFisher), ascorbic acid (50 μg/ml, Sigma), monothioglycerol (50 μg/ml, Sigma) and with different cytokines depending on the protocol. In the original VIC induction protocol, sorted day 9 CD31+/CD34+ endocardial cells were cultured for eight days in the presence of rhbFGF (10 ng/ml, R&D), rhBMP4 (100 ng/ml, R&D), TGFβ2 (0.3 ng/ml, R&D), as described in our previous work ^9^. For generation of *fibrosa*-like and *spongiosa*-like VICs with the newly established protocols, day 9 CD31+/CD34+-sorted endocardial cells were first cultured for 4 days in the presence of rhbFGF (10 ng/ml, R&D), rhBMP4 (100 ng/ml, R&D), CHIR 99021 (1 mM, Tocris) and SB-431542 (5.4 mM, Tocris)[expansion phase], followed by eight days in the presence of either rhbFGF (10 ng/ml, R&D), rhBMP4 (100 ng/ml, R&D), TGFβ2 (0.3 ng/ml, R&D)[*fibrosa* VIC specification phase] or rhbFGF (1 ng/ml, R&D) [*spongiosa* VIC specification phase].

### Monolayer differentiation of iPSCs into endocardial cells

Prior to the initiation of differentiation, iPSCs were passaged onto Matrigel coated 24-well plates at a ratio of 1:12. Differentiation was started once iPSCs reached ∼80% confluency, as previously described ^32^. Briefly, iPSC differentiation was initiated with RPMI minus insulin (RPMI 1640, supplemented with B27 minus insulin supplement (0.5x, Gibco), penicillin/streptomycin (1%, ThermoFisher), GlutaMAX (2 mM, Gibco), ascorbic acid (50 μg/mL, Sigma), apo-transferrin (150 μg/mL, R&D Systems), and monothioglycerol (50 μg/mL, Sigma). For the first 48 hours, iPSCs underwent CHIR treatment (5 µM, Tocris). From day 2 until day 6, iPSCs were treated with RPMI minus insulin supplemented with bFGF (50 ng/mL, R&D) and BMP10 (10 ng/mL, R&D). On Day 6, the differentiation medium was switched to RPMI plus insulin (RPMI 1640, supplemented with B27 supplement (0.5x, Gibco), penicillin/streptomycin (1%, ThermoFisher), GlutaMAX (2 mM, Gibco), ascorbic acid (50 μg/mL, Sigma), apo-transferrin (150 μg/mL, R&D Systems), and monothioglycerol (50 μg/mL, Sigma). From Day 6 to 14, RPMI plus insulin was further supplemented with bFGF (50 ng/mL, R&D) and BMP10 (10 ng/mL, R&D). Media was changed every other day. Endocardial differentiation was performed in a hypoxic incubator (37°C, 5% CO_2_, 5% O_2_, 90% N_2_).

### Generation of *fibrosa-*like and *spongiosa-*like VIC cells from iPSC-derived endocardial cells

On Day 14, sorted CD31^+^ endocardial cells were plated at 3×10^5^ cells per well on Matrigel-coated 12-well plates (1.25% v/v, Corning) and treated with the expansion phase medium (RPMI plus insulin supplemented with rhBMP4 [100 ng/mL, R&D], rhbFGF [10 ng/mL, R&D], CHIR [1 µM, Tocris], and SB-431542 [5.4 µM, Sigma]). After four days, the cells were treated with Phase 2 EndoMT media towards either *fibrosa* or *spongiosa* VICs for an additional eight days. *Fibrosa* VIC EndoMT was induced in RPMI plus insulin supplemented with rhBMP4 (100 ng/mL, R&D), TGFβ2 (0.3 ng/mL, R&D), and rhbFGF (10 ng/mL, R&D). *Spongiosa-*like VICs were induced with RPMI plus insulin supplemented with rhbFGF (1 ng/mL, R&D). EndoMT was performed in a normoxic incubator (37 °C, 5% CO_2_).

### Flow cytometry analyses and FACS of populations generated from HES3-NKX2-5eGFP/w and HES2:WT hESC lines

Day 3 EBs were dissociated to single cells by treatment with TrypLE for 3 minutes at 37 °C. The resulting cell suspension was filtered and transferred to IMDM medium for staining. Day 9 endocardial/endothelial monolayer cultures and VIC monolayer cultures were dissociated by incubation in collagenase type 2 (1 mg/ml, Worthington) in HANKs buffer (Gibco) at 37°C for one hour. The filtered cells were then transferred to IMDM for staining. The following antibodies were used for staining: anti-CD31-PE (BD Biosciences, 1:100), anti-CD31-AF647 (BD Biosciences, 1:100), anti-CD34-PeCy7 (eBioscience, 1:100), anti-PDGFRΒ-BV421 (BD Biosciences, 1:100), anti-CD49b-PE (BD Biosciences, 1:2000). Detailed antibody information is described in the Key Resources Table (**Table 1**). For cell-surface marker analyses, dissociated live cells were stained for 20 min at room temperature in FACS buffer consisting of PBS^-/-^ (Gibco) with 5% fetal calf serum (FCS) (Wisent) and 0.02% sodium azide. The cells were then washed twice before being subjected to further analyses. After the washing steps, stained cells were analyzed using an LSR II Flow cytometer (BD). For cell sorting, cells were stained, washed and kept in IMDM with 0.2% KnockOut^TM^ serum replacement and sorted using either a FACS Aria Fusion (BD) or FACS Symphony S6 (BD) cells sorter at the Sickids/UHN flow cytometry facility. Data were analyzed using FlowJo software (Tree Star).

**Table 1:** Key reagents and resources.

| REAGENT or RESOURCE | SOURCE | IDENTIFIER |
| --- | --- | --- |
| **Antibodies** |  |  |
| Mouse monoclonal to CD31 (clone WM59), APC conjugated | ThermoFisher | Cat.# 17-0319-42; RRID: AB_10852842 |
| Mouse monoclonal to CD31 (clone WM59), PE conjugated | BD Biosciences | Cat.# 555446; RRID: AB_395839 |
| Mouse monoclonal to CD31 (clone WM59), Alexa Fluor 647 conjugated | BD Biosciences | Cat.# 561654; RRID: AB_10896969 |
| Mouse monoclonal to CD34 (clone 4H11), PeCy7 conjugated | ThermoFisher | Cat.# 25-0349-42; RRID: AB_1963576 |
| Mouse monoclonal to CD140b (PDGFRβ) (clone 28D4), BV421 conjugated | BD Biosciences | Cat.# 564124; RRID: AB_2738609 |
| Mouse monoclonal to CD140b (PDGFRβ) (clone 28D4), BV786 conjugated | BD Biosciences | Cat.# 743038; RRID: AB_2741234 |
| Mouse monoclonal to CD49b (ITGA2) (clone 12F1), PE conjugated | BD Biosciences | Cat.# 555669; RRID: AB_396022 |
| Rabbit monoclonal to versican (VCAN) (EPR12277), non-conjugated | Abcam | Cat.# ab177480; RRID: AB_2827732 |
| Rabbit polyclonal to collagen 1 (COL1), non-conjugated | Abcam | Cat.# ab34710; RRID: AB_731684 |
| Rabbit polyclonal to collagen 3 (COL3), non-conjugated | Abcam | Cat.# ab7778; RRID: AB_306066 |
| Rabbit monoclonal to phospho-p44/42 MAPK (ERK1/2) (Thr202/Tyr204), non-conjugated | Cell Signaling Technology | Cat.# 4370s; RRID: AB_2315112 |
| Mouse monoclonal to p44/42 MAPK (ERK1/2) (L34F12), non-conjugated | Cell Signaling Technology | Cat.# 4696s; RRID: AB_390780 |
| Goat anti-rabbit IgG (H+L), HRP conjugated | Cell Signaling Technology | Cat.# 7074s; RRID: AB_2099233 |
| Horse anti-mouse IgG (H+L), HRP conjugated | Cell Signaling Technology | Cat.# 7076s; RRID: AB_330924 |
| Donkey anti-rabbit IgG (H+L), AlexaFluor488 conjugated | ThermoFisher | Cat.# A21206; RRID: AB_2535792 |
| Donkey anti-mouse IgG (H+L), AlexaFluor555 conjugated | ThermoFisher | Cat.# A31570; RRID: AB_2536180 |
| **Biological Samples** |  |  |
| HUVEC | Lonza | Cat.# C125AS |
| Primary human fetal heart tissue | This study | N/A |
| **Chemicals, Peptides, and Recombinant Proteins** |  |  |
| Penicillin/streptomycin | ThermoFisher | Cat.# 15070063 |
| L-glutamine | ThermoFisher | Cat.# 25030081 |
| GlutaMAX | ThermoFisher | Cat. #35050061 |
| Non-essential amino acids | ThermoFisher | Cat.# 11140-050 |
| Transferrin | ROCHE | Cat.# 10652202001 |
| Apo-Transferrin | R&D | Cat.# 3188-AT |
| Ascorbic acid | Sigma | Cat.# A4544-100G |
| Monothioglycerol | Sigma | Cat.# M-6145-25ml |
| β-Mercaptoethanol | ThermoFisher | Cat.# 21985-023 |
| ROCK inhibitor Y-27632 | Tocris | Cat.# 1254 |
| Recombinant human BMP4 | R&D | Cat.# 314-BP |
| Recombinant human ActivinA | R&D | Cat.# 338-AC |
| Recombinant human TGFβ2 | R&D | Cat.# 302-B2 |
| Recombinant human bFGF | R&D | Cat.# 233-FB |
| Recombinant human FGF4 | R&D | Cat.# 235-F4 |
| Recombinant human BMP10 | R&D | Cat.# 2926-BP |
| Recombinant human VEGF | R&D | Cat.# 293-VE |
| IWP2 (Wnt inhibitor) | Tocris | Cat.# 3533 |
| CHIR 99021 (GSK-3 inhibitor) | Tocris | Cat.# 4423 |
| SB-431542 (TGFβ inhibitor) | Sigma-Aldrich | Cat.# S4317 |
| Collagenase type2 | Worthington | Cat.# LS004-176 |
| DAPI | Biotium | Cat.# 40043 |
| Fetal bovine serum (FBS) | ThermoFisher | Cat.# 16000044 |
| Fetal calf serum (FCS) | Wisent | Cat.# 080-150 |
| Sodium azide | Millipore Sigma | Cat.# S2002 |
| Bovine serum albumin (BSA) | Sigma | Cat.# A1470 |
| Matrigel, growth factor reduced | Corning | Cat.# 356230 |
| Accutase | Innovative Cell Technologies, Inc | Cat.# AT104 |
| Thiazovivin | Selleckchem | Cat.# S1459 |
| DNase 1 | Millipore Sigma | Cat.# 260913 |
| HEPES | Research Products International | Cat.# H75050 |
| Triton X-100 | MilliporeSigma | Cat.# 93418 |
| RIPA buffer | MilliporeSigma | Cat.# R0278 |
| Laemmli Sample Buffer | Bio-Rad | Cat.# 1610747 |
| Tris/Glycine/SDS Buffer | Bio-Rad | Cat.# 1610732 |
| Amersham ECL Prime Western Blotting Detection Reagent | Cytiva | Cat.# RPN2236 |
| Restore PLUS Western Blot Stripping Buffer | ThermoFisher | Cat.# 46430 |
| Halt Protease and Phosphatase Inhibitor | ThermoFisher | Cat.# 78440 |
| Fluo-4 | ThermoFisher | Cat.# F14201 |
| PPADS tetrasodium salt | Tocris | Cat.# 0625 |
| Shark cartilage chondroitin sulfate | MilliporeSigma | Cat.# C4384 |
| Hydroxyproline | MilliporeSigma | Cat.# PHR1939 |
| **Critical Commercial Assays** |  |  |
| RNAqueous-micro kit with RNase-free Dnase treatment | Ambion | Cat.# AM1931 |
| iSCRIPT Reverse Transcriptase | Bio-Rad | Cat.# 1798841 |
| QuantiFast SYBR Green PCR kit | QIAGEN | Cat.# 204057 |
| CD31 MicroBead Kit, human | Miltenyi Biotec | Cat.# 130-091-935 |
| CD34 MicroBead Kit, human | Miltenyi Biotec | Cat.# 130-046-702 |
| MACS cell separation, MS columns | Miltenyi Biotec | Cat.# 130-042-201 |
| MACS cell separation, LS columns | Miltenyi Biotec | Cat.# 130-042-401 |
| Pierce BCA Protein Assay Kit | ThermoFisher | Cat.# 23225 |
| 10x Genomics Single Cell 3' v3.1 Reagent Kit | 10x Genomics | Single Cell 3' v3.1 |
| **Deposited Data** |  |  |
| Raw data files for primary human fetal heart scRNA-seq | This study | GEO: GSE228638 |
| Raw data files for hPSC-derived endocardial/endothelial scRNA-seq | 9 | GEO: GSE151186 |
| Raw data files for hPSC-derived VICs scRNA-seq | This study | GEO: GSE234382 |
| **Experimental models: Cell Lines** |  |  |
| Human ESC: HES3-NKX2-5<sup>eGFP/w</sup> line | Gift from Drs. E. Stanley and A. Elefanty, Monash University, AU <sup>48</sup> | N/A |
| Human ESC: HES2 line | WiCell | Cat.# ES02 |
| Human iPSC: WTC11 line (XY, mono-allelic ACTN2-mEGFP) | Coriell Institute | #AICS-0075-085 |
| Human CRISPR *PTPN11*<sup>N308D/N308D</sup> iPSC line | Synthego | N/A |
| **Oligonucleotides** |  |  |
| See Table 2-2 for PCR primer sequences | This study | Table 2-2 |
| **Software and Algorithms** |  |  |
| FlowJo | Tree Star | https://www.flowjo.com |
| ImageJ | FIJI | https://imagej.nih.gov/ij/ |
| GraphPad Prism | GraphPad Software | http://www.graphpad.com/scientific-software/prism |
| Metamorph v7.8.2.0 | Molecular Devices LLC |  |
| Pannoramic Viewer 1.15.4 | 3DHistech | https://www.3dhistech.com/research/software-downloads/ |
| 10x Chromium Single Cell Software Suite v3.1 | 10x Genomics | https://support.10xgenomics.com/single-cell-gene-expression/software/pipelines/latest/what-is-cell-ranger |
| R Version 3.6.1 |  | https://cran.r-project.org/ |
| Seurat, Version 4.3.0 | <sup>55, 56</sup> | https://github.com/satijalab/seurat |
| Bioconductor v3.10 |  | https://bioconductor.org/news/bioc_3_10_release/ |
| Cell Ranger v2.1.0 | 10x Genomics | https://support.10xgenomics.com/single-cell-gene-expression/software/pipelines/latest/what-is-cell-ranger |
| Bader Lab Gene Sets |  | http://baderlab.org/GeneSets |
| SingScore v1.16 | <sup>24</sup> | https://github.com/DavisLaboratory/singscore |
| **Other** |  |  |
| StemPro-34 media (kit) | ThermoFisher | Cat.# 10639011 |
| DMEM, high glucose | ThermoFisher | Cat.# 11965092 |
| DMEM/F12 | Cellgro | Cat.# 10-092-CV |
| RPMI 1640 | ThermoFisher | Cat.#11875093 |
| B27 Supplement | ThermoFisher | Cat.#17504044 |
| B27 Supplement, Minus Insulin | ThermoFisher | Cat.#A1895601 |
| EBM-2 (Endothelial Basal Medium) | Lonza | Cat# CC-3156 |
| EGM-2 (SingleQuots supplement kit for EBM-2) | Lonza | Cat# CC-4176 |
| PBS no Mg, no Ca | ThermoFisher | Cat.#14190-250 |
| Hank's Balanced Salt Solution | ThermoFisher | Cat.# 14175-079 |
| KnockOut serum replacement | ThermoFisher | Cat.# 10828028 |
| mTesR Plus | Stem Cell Technologies | Cat.# 100-1130 |
| TrypLE | ThermoFisher | Cat.# 12605010 |
| 6-well clear flat bottom TC-treated culture plate | Corning | Cat.# 353046 |
| 12-well clear flat bottom TC-treated culture plate | Corning | Cat.# 353043 |

### Flow cytometry analyses and FACS of populations generated from iPSC lines

Endocardial cells and VICs were dissociated with collagenase II (0.6 mg/ml, Worthington) in Hank’s buffer (Gibco) supplemented with HEPES (5 mM, Sigma), incubated at 37 °C for 1 hour. Dissociated endocardial cells were collected, stained with CD31 microbeads (Miltenyi) and sorted to >95% purity on LS Columns. VICs were similarly stained with CD31 microbeads and negative selection for CD31^-^ population was performed on LD columns. Flow cytometry analysis of these populations was performed following dissociation. Antibody staining was performed in FACS staining buffer, consisting of PBS^-/-^ (Gibco) supplemented with FBS (10%, Gibco), DNase (10 μg/mL, Millipore), and thiazovivin (2 μM, Selleckchem), on ice for one hour, followed by wash steps with PBS. Samples were resuspended in FACS running buffer consisting of PBS^-/-^ (Gibco) supplemented with FBS (1%, Gibco), penicillin/streptomycin (2%, ThermoFisher), DNase (10 μg/mL, Millipore), thiazovivin (2 μM, Selleckchem), and DAPI (0.1 μg/mL, Sigma) and analyzed on either BD FACS Celesta, BD LSR Fortessa, or BD LSR II flow cytometers. Antibodies used were anti-CD31 APC (1:100, ThermoFisher) and anti-PDGFRβ BV786 (1:100, BD Biosciences).

### Immunohistochemistry of cells generated from HES2:WT hESC lines

CD34^+^ endocardial cells generated from HES2:WT line were plated onto 12-mm cover glasses (VWR) pre-coated with Matrigel (25% v/v, BD) in 24-well plates (Corning) at a density 2×10^5^ cells per well. The cells were cultured for 12 days as monolayers under *fibrosa* VIC or *spongiosa* VIC differentiation conditions. Following culture, the cells were fixed with 4% paraformaldehyde in PBS for 10 min at room temperature and subsequently permeabilized with PBS containing 0.2% TritonX for 20 min. The fixed cells were blocked with PBS containing 10% FCS and 2% BSA. The following antibodies were used for staining: rabbit anti-Versican (Abcam, 1:100), mouse anti-human CD49b (BD Biosciences, 1:100). For detecting unconjugated primary antibodies, the following secondary antibodies were used: donkey anti-mouse IgG-A555 (ThermoFisher, 1:1000), donkey anti-rabbit IgG-A488 (ThermoFisher, 1:1000). Detailed antibody information is described in the Key Resources Table (**Table 1**). The cells were stained with primary antibodies in staining buffer consisting of PBS with 0.05% TritonX and 2% BSA overnight at 4 °C. The stained cells were washed with PBS containing 0.1% BSA 3x for 10 min each wash at room temperature. The cells were then stained with secondary antibodies in staining buffer for one hour at room temperature followed by a wash step as described above. The cell nuclei were stained with DAPI (Biotium, 0.3 μg/ml) in wash buffer for 5 min at room temperature. Following staining, the samples were mounted using ProLong Diamond Antifade Mountant (ThermoFisher). All images were captured using a Zeiss LSM700 confocal microscope and analyzed using Image J software (NIH).

### Human tissue-engineered extracellular matrix (hTEM) production

Non-woven polyglycolic acid (PGA) mesh (SaniSure SA) was cut into 6-cm^2^ circles and coated with 1% poly-4-hydroxybutyrate (P4HB) (TEPHA, Inc) dissolved in liquid tetrahydrofuran (Sigma). Polymeric scaffolds were then sutured onto stainless steel rings and sterilized in 70% ethanol, as previously described ^51^. hPSC-derived VICs (derived from either the *fibrosa* or *spongiosa* protocol) were seeded at a 1×10^6^ cells/cm^2^ density onto scaffolds using fibrin as a cell carrier ^52^. Seeded scaffolds were then cultured on an orbital shaker at 37 °C for four weeks. StemPro medium was supplemented with 0.25 mg/mL L-ascorbic acid two-phosphate (Sigma) as well as 0.3 ng/mL or 1.0 ng/mL TGFβ2 (R&D Systems), as indicated. Following four weeks of culture, hTEMs were decellularized with a detergent-based solution containing 0.25% Triton X-100, sodium deoxycholate, and 0.02% EDTA, as previously described ^44^.

### hTEM histology and immunohistochemistry

Qualitative measurements of hTEM matrix protein content was demonstrated using histology and immunohistochemistry. hTEMs were fixed in 10% formalin, paraffin-embedded, and consecutively cut into 5-μm sections. Sections were analyzed using H&E, collagen 1 (Abcam, ab34710; 1:200), collagen 3 (Abcam, ab7778; 1:250), versican (Abcam, ab177480, 1:50), and picrosirius red. Samples were imaged using a brightfield slide scanner (Mirax Midi Microscope, Carl Zeiss GMbH) and analyzed using Pannoramic Viewer software (3DHistech). Polarized light microscopy was performed at the VHX-970 System (Keyence).

### Biochemical analyses of hTEMs

Glycosaminoglycan and hydroxyproline, an indicator of collagen content, were quantified using previously described protocols ^53, 54^. Briefly, 2-3 mg of lyophilized hTEMs were digested in a 6-mM papain (Sigma) solution at 60 °C overnight. Glycosaminoglycan content was compared to shark cartilage chondroitin sulfate (10 mg/mL, Sigma) using a colorimetric dimethylmethylene blue (Sigma) solution ^53^. Hydroxyproline content was compared to a 10 mg/mL hydroxyproline standard (Sigma) using an aldehyde/perchloric acid solution, as previously described ^54^. All measurements were performed in triplicate and concentrations were compared to the total dry weight of each hTEM.

### FGF4 stimulation and western blot analysis

Following CD31 negative selection for *fibrosa-*like VICs, VICs were plated on Matrigel coated 6-well plates (1.25% v/v, Corning) in RPMI plus insulin supplemented with FBS (5%, Gibco) and rhbFGF (10 ng/mL R&D). VICs were allowed to grow to confluency, at which point they were serum starved overnight in DMEM (Gibco) supplemented with FBS (0.1%, Gibco). Following serum starvation, media was replaced with DMEM (Gibco) supplemented with FGF4 (25 ng/mL, R&D) for the indicated time points. VICs were then washed with PBS and lysed on ice with RIPA (Sigma) + Halt Protease and Phosphatase Inhibitor (ThermoFisher). Protein concentration was determined with Pierce Protein BCA Assay (ThermoFisher).

For western blotting, 15 µg of total protein was boiled at 95 °C for five minutes in Laemmlli sample buffer (BioRad). SDS-PAGE was then performed on a 7.5% Mini-PROTEAN TGX gel (BioRad) in Tris/Glycine/SDS buffer (BioRad). The protein was then transferred to a nitrocellulose membrane (BioRad) in Tris/glycine buffer (BioRad) with 20% methanol (Sigma). Membranes were blocked in blocking buffer containing 5% BSA (Sigma) in TBST (QuickSilver). Primary antibody incubation for either rabbit anti-pERK (Cell Signaling Technology #4370S, 1:1000) or mouse anti-total ERK (Cell Signaling Technology #4696S, 1:1000) was performed overnight at 4 °C on shaker in blocking buffer. Secondary antibody incubation with either anti-rabbit HRP (Cell Signaling Technology #7074S, 1:15,000) or anti-mouse HRP (Cell Signaling Technology #7076S, 1:15,000) was performed in blocking buffer for one hour at room temperature. Detection of chemiluminescent signal was performed with ECL Prime Western Blotting Detection Reagent (Amersham) and detected with BioRad ChemiDoc. Membranes were stripped with Restore PLUS Western Blot Stripping Buffer (ThermoFisher), after which membranes were blocked and incubated with antibodies as described above. Densitometry analysis of blots was performed with FIJI software.

### Calcium transient measurements

Day 7 endocardial and control endothelial cultures generated from HES2:WT hPSC line were dissociated and the CD31^+^PDGFRβ^+^ and CD31^+^PDGFRΒ^-^ endocardial populations and CD31^+^PDGFRβ^-^ control endothelial populations isolated by FACS. The sorted cells, as well as HUVECs, were then plated in a 12-well flat bottom microplate (Corning) coated with Matrigel (25% v/v, Corning) at a density of 5×10^5^ cells per well and cultured for two days. Endocardial and control endothelial cells were cultured in StemPro-34 supplemented with penicillin/streptomycin (1%, ThermoFisher), L-glutamine (2 mM, ThermoFisher), ascorbic acid (50 μg/ml, Sigma), monothioglycerol (50 μg/ml, Sigma), transferrin (150 μg/ml, ROCHE), rhbFGF (50 ng/ml, R&D) and with either rhBMP10 (10 ng/ml)[endocardium] or VEGF (100 ng/mL, R&D)[control endothelium]. HUVECs were cultured in EGM-2 medium (Lonza). The cultures were incubated in a low oxygen environment (5% CO_2_, 5% O_2_, 90% N_2_).

On day 9, the monolayers were incubated with Fluo-4 (Invitrogen, final concentration; 4 uM) for 30 min at 37 °C. The monolayers were then washed once with PBS^-/-^ and subsequently incubated in PBS^-/-^ for an additional 10 minutes, with or without the presence of the nonselective P2X antagonist PPADS (Tocris, final concentration; 3 uM). The monolayers were imaged using a Zoom microscope body MVX10 with the objective MVPLAPO 0.63X (NA 0.15, WD 87 mm, FN 22, Olympus) for an overall FOV of 10 mm x 10 mm with the wavelength 490–535 nm for the excitation and 532–588 nm for bandpass-filter Fluo-4. Five seconds after the start of recording, ATP (Sigma, final concentration; 10 uM) was added to the monolayers and calcium transients were measured using MetaMorph software.

### Quantitative reverse transcription polymerase chain reaction (RT-qPCR)

Total RNA extraction was performed using the RNAqueous-micro Kit (Invitrogen). Up to 1 ug of RNA was treated with RNase-free DNase (Invitrogen) and then reverse transcribed into cDNA using oligo (dT) primers and random hexamers and iSCRIPT reverse transcriptase (ThermoFisher). RT-qPCR was performed on an EP Real-Plex MasterCycler (Eppendorf) using the QuantiFast SYBR Green PCR kit (QIAGEN). All data were generated as technical duplicates evaluated for relative copy number, reaction efficiency, and genomic DNA contamination (< 0.01% of TBP content) using a 10-fold dilution series of sonicated human genomic DNA standards made in house from wild-type HES2 hESC cells ranging from 25 ng/ml to 2.5 pg/ml. Samples free of genomic DNA contamination were assessed for expression of genes of interest relative to the house keeping gene *TBP*. The primer sequences are provided in **Table 2**.

**Table 2:**
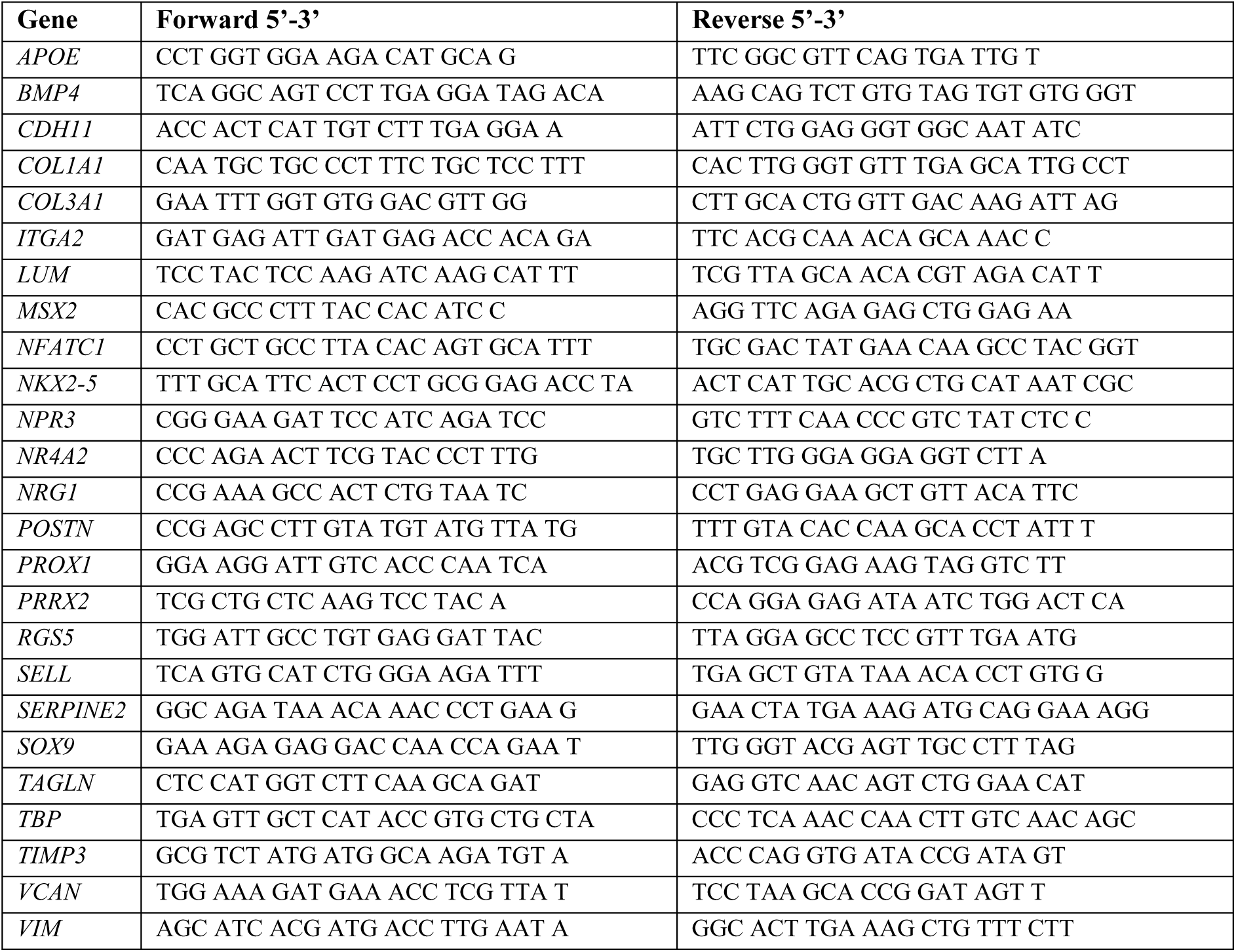
RT-qPCR primer sequences.

### Sample preparation and single-cell library generation of human fetal heart tissue

De-identified human fetal heart tissues were obtained in collaboration with the University of Washington Birth Defects Research Lab (BDRL, IRB STUDY00000380), University of Southern California and Children’s Hospital Los Angeles, and Cincinnati Children’s Research Foundation (CCRF, IRB 2011-2856). All samples were collected from parents who provided signed informed consent according to institutional review board (IRB) guidelines in accordance with all university, state, and federal regulations. The only clinical information collected was gestational age and the presence of any maternal or fetal diagnoses. Fetal tissues for scRNA-seq and primary cell culture were transferred from BDRL to Stanford University under Material Transfer Agreement #44556A. The samples, data, and/or services derived from CCRF were provided by Discover Together Biobank. The above studies were classified as exempt because of the de-identified nature of the valve samples.

One normal human fetal heart (week 15, male) was manually micro-dissected. The heart was kept on ice throughout processing and 1X HBSS (GIBCO) was used to remove excess blood cells. Aortic, pulmonary, mitral, and tricuspid valves were collected. Digestion buffer was prepared in DMEM medium and included the following components: Liberase TM, 0.5 mg/ml; DNase I, 20 μg/ml; HEPEs, 10 mM. Following digestion for 10 min at 37 °C, EGM-2 medium was added to stop the reaction and cells were passed through a 40-μm cell strainer. Populations with viability > 90% were sent to Stanford Functional Genomics Facility (SFGF) for 10X chromium single-cell RNA-seq paired-end library preparation using V2 version (10X Genomics). All samples were uniquely indexed, mixed, and evenly distributed into the Illumina HiSeq 4000 for sequencing. The raw and processed data from single-cell RNA sequencing in this study have been deposited with the Gene Expression Omnibus (https://www.ncbi.nlm.nih.gov/geo/) under accession number GSE228638.

### Sample preparation and single-cell library generation of hPSC-derived endocardium

scRNA-seq data associated with hPSC-derived endocardial cells have been previously published by our group ^9^ and are deposited in GEO (GSE151186). Briefly, putative endocardial and control endothelial cells were generated from the HES2:WT hESC line ^49^, dissociated to single cells at day 9, and stained with DAPI to exclude dying cells. Live cells were sorted using FACSAria Fusion (BD) at the Sickkids/UHN flow cytometry facility. After live cell sorting, the endocardial bulk population was stained with TotalSeq-A0255 anti-human Hashtag 5 anti-body (BioLegend), while the control endothelial bulk populations were stained with TotalSeq-A0254 anti-human Hashtag 4. The two populations were then mixed and single-cell libraries were generated in the 10X Genomics Chromium controller using the Chromium Single Cell 3′ Reagent Kit v3. Hashtagging facilitated the differentiation between hPSC-derived endocardial and control endothelial cells following sequencing.

### Sample preparation and single-cell library generation of hPSC-derived VICs

Putative *fibrosa* and *spongiosa* VICs were generated from the HES2:WT hESC line ^49^, dissociated to single cells at day 9+12, and stained with DAPI to exclude dying cells. Live cells were sorted using FACSAria Fusion (BD) at the Sickkids/UHN flow cytometry facility. After live cell sorting, *fibrosa* and *spongiosa* VIC populations were labeled with the 10X Genomics 3’ CellPlex Kit. The two populations were then mixed and single-cell libraries were generated in the 10X Genomics Chromium controller using the Chromium Single Cell 3′ Reagent Kit v3. Multiplexing facilitated the differentiation between hPSC-derived *fibrosa*-like and *spongiosa*-like VICs following sequencing. Sequencing data are deposited in GEO (GSE234382).

### scRNA-seq analyses

Chromium Single Cell Software Suite v3.1 was used for processing the scRNA-seq data produced in the 10x Chromium Platform. The ‘‘cellranger mkfastq’’ pipeline was used to demultiplex raw base call (BCL) files into FASTQ files. Reads in the FASTQ files were then mapped to the human reference genome (NCBI build38/UCSC hg38) using the STAR software. Chromium cellular barcodes were then used to generate gene-barcode matrices. Only reads confidently (uniquely) mapped to the transcriptome were used for the UMI count. Filtered gene-barcode matrices containing only cellular barcodes were used for downstream analysis.

The R-based (R 3.6.1) package Seurat v4.3.0 was utilized for all downstream single cell analyses ^55, 56^. Data were further filtered by excluding empty droplets and low-quality cells that expressed less than 500 genes, as described in published tutorials (http://satijalab.org/seurat/). Putative dead or lysed cells were excluded by removing cells in which a high percentage of transcripts mapped to mitochondrial genes. Ultimately, the downstream analyses encompassed a total of 58,996 primary fetal heart cells, 3,965 hPSC-derived endocardial cells, and 5,953 hPSC-derived VICs.

Seurat’s SCTransform v2 pipeline ^57, 58^ was used for data normalization and scaling and to identify highly variable features. The variance-stabilizing transformation (*VST*) method in SCTransform was used to select the top 3000 highly variable genes. Subsequently, principal component analysis (PCA) was performed for dimensionality reduction (*RunPCA* function). For each dataset, the top 25–50 principal components were selected based on the *ElbowPlot* method and subsequently used to construct the nearest neighbor graph (*FindNeighbors* function). This information was then used for downstream graph-based clustering (*FindClusters* function) using the Louvain algorithm and a resolution parameter of 0.4–1.2. Finally, Uniform Manifold Approximation and Projection (UMAP) was used for non-linear dimensionality reduction and projection to a 2D plot for visualization using the selected top principle components.

For primary human fetal heart cells, clustering was performed by using the top 50 principal components and by setting the resolution parameter to 0.4. Fetal heart cell types were annotated based on canonical cell type markers in the human heart. Irrelevant cell types, including leukocytes, erythrocytes, platelets, pericardium, and an outlier cluster with high ribosomal protein expression were removed. A subset of 20,889 VICs were extracted from the fetal heart data and were analyzed using the standard pipeline described above. The top 30 principal components were used for clustering, and the clustering resolution parameter was set to 0.8. Clusters expressing *SCX*, *MXRA5*, and *TIMP3* were designated as *fibrosa* VIC clusters, whereas clusters expressing *LUM* and *OGN* were designated as *spongiosa* VIC clusters. For hPSC-derived endocardial cells, clustering was performed by using the top 25 principal components and by setting the resolution parameter to 1.2. For hPSC-derived VICs, clustering was performed by using the top 50 principal components and by setting the clustering resolution parameter to 0.9.

Differential expression analyses between specific clusters were performed using Seurat’s built-in *FindMarkers* function. DEGs were computed using the Wilcoxon rank sum test (min.pct: 0.1; logFC threshold: 0.25; adjusted p-value < 0.05). Gene set enrichment analyses (GSEA) were performed using the *fgsea* package v1.22.0. For our GSEA analyses, we used the Bader Lab Gene Sets, updated March 2022 (http://baderlab.org/GeneSets). These gene sets have been assembled by the Bader Lab at the University of Toronto and provide a more focused and curated selection of gene sets derived from the broader Gene Ontology database.

Mapping of hPSC-derived VICs to reference primary fetal VICs was performed using the *FindTransferAnchors* and *MapQuery* functions in Seurat. The *FindTransferAnchors* function identifies a set of anchor features that capture the shared biological signal between the reference and query objects. An anchor feature is a gene or a combination of genes that exhibit similar expression patterns across different datasets. Once the anchor features are identified using *FindTransferAnchors*, the *MapQuery* function is used to project/query new cells or datasets onto the integrated reference space. This function maps or aligns new scRNA-seq data onto the reference dataset, allowing for the comparison and analysis of cells from different datasets in a common integrated space.

To measure the similarity of hPSC-derived VICs to primary fetal heart cells, we used correlation analysis. To mitigate the impact of drop-out events inherent to scRNA-seq, we first created pseudo-bulks by aggregating the expression profiles of similarly annotated cells in the primary fetal heart dataset and of hPSC-derived VICs. Correlation between hPSC-derived VICs and human fetal cardiac cells was estimated with Spearman’s correlation based on the mean expression values of commonly expressed genes.

To further assess the similarity of hPSC-derived VICs to the various subpopulations present in the fetal heart, we used the SingScore v1.16 ^24^ package for gene set scoring. Compared with other scoring approaches, such as single-sample GSEA, the SingScore method demonstrates a high level of consistency across independent data sets, resulting in easily interpretable scores. We first identified the top 100 significantly (adjusted p-value < 0.01) up- or down-regulated markers specific to each fetal subpopulation. We then applied the SingScore package to measure the aggregate expression score of these identified fetal subpopulation markers within the hPSC-derived VICs.

## Sources of funding

Canadian Institutes of Health Research (FDN159937)

Canadian Institutes of Health Research Fellowship Award

Thoracic Surgery Foundation Resident Research Fellowship Award

American Association for Thoracic Surgery Cardiac Surgical Resident Investigator Award

Mäxi Foundation

European Research Council (ERC) grant agreement no. 852814 (TAVI4Life)

## Disclosures

G. M. K. is a founding investigator and a paid consultant for BlueRock Therapeutics LP and a paid consultant for VistaGen Therapeutics. S.P.H. is a shareholder at Xeltis BV and LifeMatrix Technologies AG. M.Y.E. is a shareholder at LifeMatrix Technologies AG.

## Resource tables

**Supplementary Table 1:** Differentially expressed genes (DEGs) between *Fibrosa* and *Spongiosa* valvular interstitial cells (VICs) in the primary human fetal valve scRNA-seq dataset.

**Supplementary Table 2:** Differentially expressed genes (DEGs) between *Fibrosa* and *Spongiosa* valvular interstitial cells (VICs) in the hPSC-derived VIC scRNA-seq dataset.

## Supplementary figure legends

**Supplementary Figure 1. Expression of ACTA2 in endocardial-derived mesenchymal cells cultured on substrates of different stiffness**

Immunocytochemistry for α-smooth muscle actin (ACTA2, red) and nuclei (DAPI, blue) in hPSC-derived VICs cultured on soft (5 kPa, left) and stiff (1.72 MPa, right) substrates. Cells on the stiff substrate exhibit robust ACTA2 expression and organized stress fibers, indicative of myofibroblastic activation. Scale bar = 100 µm.

**Supplementary Figure 2. Effect of bFGF, BMP4 and TGFβ2 on generation of VICs from SB/CHIR treated endocardial cells**

**(A)** Schematic of the experimental strategy used to analyze the effects bFGF, BMP4 and TGFβ2 in the specification phase of the new VIC protocol. **(B)** Representative flow cytometric analyses of CD31 and PDGFRβ expression on VICs generated at day 12 following culture in the indicated factors during the specification phase (day 4–12) using the protocol described in (A). **(C)** Quantification of the total number of CD31^-^PDGFRβ^+^ hPSC-derived VICs generated at day 12 following culture in the indicated factors during the specification phase (day 4–12). (n = 6, one-way ANOVA with Tukey’s multiple comparisons test, *p < 0.05, **p < 0.01,***p < 0.001; error bars ± SEM). **(D)** RT-qPCR analysis of the expression levels of VIC-specific genes (*PRRX2*, *TIMP3*, *ITGA2*, *NR4A2*) in CD31^-^PDGFRβ^+^ hPSC-derived VICs at Day9 +12 generated with the indicated factors during the specification phase in the protocol in (A). (n = 3, one-way ANOVA with Tukey’s multiple comparisons test, *p < 0.05, **p < 0.01,***p < 0.001; error bars ± SEM). For all RT-qPCR analyses, expression values were normalized to the housekeeping gene *TBP*. Error bars represent SEM. *D*, day; *EndoMT*, endothelial-to-mesenchymal transition; *MACS*, magnetic-activated cell sorting; *F*, bFGF; *B*, BMP4.

**Supplementary Figure 3. VIC potential of PDGFRβ^+^ and PDGFRβ^-^ endocardial cells**

**(A)** Schematic of the protocol used to generate and isolate hPSC-derived CD31^-^PDGFRβ^+^ mesenchymal cells from PDGFRβ positive and negative NKX2-5^+^CD31^+^ endocardial cells. **(B)** Representative flow cytometric analyses of NKX2-5, CD31, and PDGFRβ expression on the day 9 endocardial population and of PDGFRβ and CD31 expression on the mesenchymal cells generated from the isolated PDGFRβ positive and negative NKX2-5^+^CD31^+^ endocardial cells.

**(C-E)** Quantification of the frequency of CD31^-^PDGFRβ^+^ cells (C), total cell numbers (D) and total number of CD31^-^PDGFRβ^+^ cells (E) generated from PDGFRβ positive and negative NKX2-5^+^CD31^+^ endocardial cells following protocol described in (A)(n = 4, paired t-test *p < 0.05, **p < 0.01, ***p < 0.005).

Error bars represent SEM. *D*, day; *EndoMT*, endothelial-to-mesenchymal transition; *FACS*, fluorescence-activated cell sorting.

**Supplementary Figure 4. Expression of CD49b in hPSC-derived *fibrosa* and *spongiosa* VICs**

**(A)** Gating strategy for the flow cytometry analysis showing CD49b and CD31 expression on the VIC population in **Fig 5C** generated with the protocol described in **Supplementary Fig 2A**. This protocol yields a heterogeneous VIC-like population containing both CD49b⁺ and CD49b⁻ cells.

**(B)** Gating strategy for flow cytometric analyses of *fibrosa* and *spongiosa* VIC populations in **Fig 5H** (generated with the protocols described in **Fig 5G**) showing the levels of CD49b expression in these populations.

**Supplementary Figure 5. Gene set analysis of hPSC-derived *fibrosa*-like and *spongiosa*-like VICs**

Gene set scoring analysis of hPSC-derived *fibrosa*-like (red) and *spongiosa*-like (blue) VIC populations, showing SingScore values for primary human fetal heart cell type signatures.

**Supplementary Figure 6. Evaluation of collagen maturity in human tissue-engineered extracellular matrices**

Representative qualitative images of hTEMs from **Fig 7C** subjected to picrosirius staining in brightfield (top row) and under polarizing light (bottom row). Control hTEMs (last column) were seeded with human dermal fibroblasts. Scale bars indicate 500 μm.

## Notes

### Competing Interest Statement

The authors have declared no competing interest.

